# Biotic interactions shape the realised niche of toxic cyanobacteria

**DOI:** 10.1101/2025.04.16.649070

**Authors:** Pinelopi Ntetsika, Stefanie Eyring, Ewa Merz, Marta Reyes, Benno Käch, Stephan Munch, Stuart Dennis, Marco Baity-Jesi, Francesco Pomati

## Abstract

Cyanobacterial blooms increasingly threaten vital freshwater ecosystems, with harmful impacts exacerbated by climate change and eutrophication. Despite extensive research on temperature and nutrient effects, our predictive capacity remains limited. We propose that this limitation stems from insufficient understanding of how biotic interactions modify cyanobacterial responses to abiotic conditions. Using five years of daily monitoring data from a eutrophic lake and state-space reconstruction modelling, we show that interactions with co-occurring plankton species fundamentally reshape the realised niche of bloom-forming cyanobacteria. Biotic interactions shift temperature thresholds by up to 13°C and phosphorus requirements by over 20 μg/L—effects substantial enough to determine whether environmental conditions support or prevent blooms in *Microcystis* and *Dolichospermum*. Grazing inhibits bloom formation across cyanobacterial taxa, while facilitation by other phytoplankton may allow blooms at unexpectedly low temperatures and phosphate concentrations. These findings address a fundamental research gap—how species interactions shape realised niches in natural systems—while offering practical insights for bloom management. By integrating biotic interactions into monitoring programs and predictive models, we can improve forecasting accuracy and develop targeted interventions that complement traditional nutrient control approaches. These findings parallel recent advances in ecology suggesting the fundamental role of biotic interactions in mediating species’ responses to environmental change.

## Main text

Cyanobacterial blooms significantly impact freshwater ecosystems, causing substantial economic losses in services like recreation, drinking water, and fisheries^1,2^. The global prevalence of these blooms increased over recent decades, with projections suggesting continued expansion^1,3^. Beyond economic impacts, cyanobacteria produce bioactive metabolites that are toxic to animals and potentially harmful to humans, particularly when biomass accumulates to high concentrations^3,4^. Despite their ecological and economic importance, predicting bloom occurrence and magnitude remains challenging. Current research on cyanobacterial blooms primarily emphasizes abiotic factors—nutrient availability (phosphorus and nitrogen) and temperature—as key drivers of growth and toxic outbreaks^3,5–7^. However, bloom dynamics result from the balance between cell growth and loss processes, including mortality and physical removal^8^. While we recognise that blooms emerge from complex networks of abiotic factors and species interactions with grazers and competitors^9^, the specific role of biotic interactions in natural systems remains poorly understood. This limitation potentially undermines our forecasting models, which may over- or underestimate bloom occurrence and magnitude when biotic factors are neglected^10^. Here, we address this critical knowledge gap—the insufficient understanding of how biotic interactions shape cyanobacterial blooms in natural systems—and hypothesise that these interactions provide the mechanistic understanding needed to explain and ultimately predict the complex ecological dynamics driving bloom formation.

While we hypothesise that biotic interactions are crucial for understanding bloom formation, current knowledge of these interactions stems primarily from controlled laboratory experiments with limited taxa and short timescales. Studies have focused predominantly on competitive and predator-prey dynamics, with less attention to facilitative mechanisms. Cyanobacteria possess well-documented competitive advantages in warm, nutrient-rich environments, including CO_2_-concentrating mechanisms, specialised pigmentation, nitrogen fixation, and buoyancy control^3^. Their bioactive metabolites may play a role in stress responses^11^ and as mediators of biotic interactions, particularly with zooplankton^12^. Cyanobacteria’s low nutritional value, toxicity, and large colony size make them unfavourable prey, with some zooplankton (e.g. copepods) promoting cyanobacterial dominance through selective grazing that targets competing algae^13^. Conversely, microzooplankton (especially rotifers) and cladocerans, may control blooms through direct consumption^14,15^. Critically, these biotic interactions are dynamic—shifting with environmental conditions^16,17^ and through zooplankton physiological and behavioural adaptations occurring on short timescales^12,14^. The research gap lies specifically in understanding how dynamic biotic interactions shape cyanobacteria’s realised niche across temperature and phosphorus gradients—the very environmental factors most affected by climate change and eutrophication.

Here, we investigate whether and how interactions between cyanobacteria and co-occurring plankton taxa influence cyanobacterial growth, particularly their net accumulation across abiotic gradients. Rather than using arbitrary abundance thresholds to define blooms, we focus on net growth rates (nGR)—where biomass accumulation exceeds loss at a daily scale. This approach captures the rapid dynamics of bloom formation and enables earlier detection by both researchers and stakeholders^8,9^. We analyse daily data from Lake Greifensee (Switzerland) across five consecutive growing seasons (1742 days), using an automated underwater microscope (see **materials and methods**) that records organisms from 10 μm to 1 cm—spanning from cyanobacteria to invertebrate predators, herbivores of various sizes, and mixotrophic and photosynthetic protists—encompassing 83 taxa (**Fig. 1** and **Tables S1** and **S2**). A neural network classifies each detected object to generate a daily time series of taxonomic abundances (**Figs. S1 to S3** and **Table S3**). Concurrently, we monitor all major physical and chemical parameters influencing cyanobacterial growth, including water temperature, nutrients, and meteorological conditions (**Fig. 1** and **Fig. S4**), with all abiotic data quality-controlled and aggregated to daily resolution (see **materials and methods**).

**Fig. 1:**
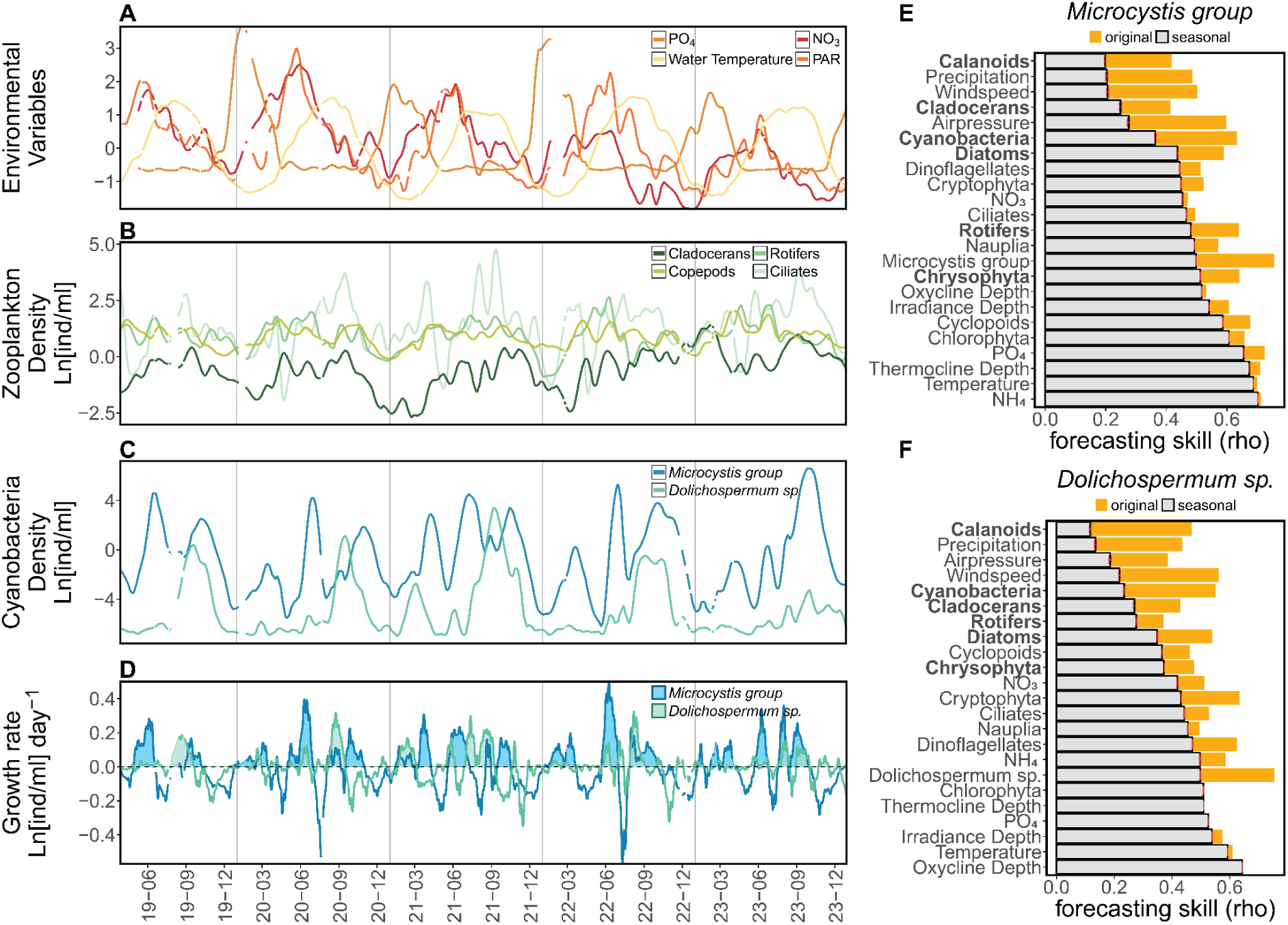
Environmental conditions and cyanobacterial population dynamics in Lake Greifensee (2019-2023). (**A**) Seasonal patterns in the temperate eutrophic lake showing scaled values of water temperature (°C), photosynthetically active radiation depth (PAR, m), phosphate (PO_4_, μg/L), and nitrate (NO_3_, mg/L): nutrients peak during winter circulation when temperatures are lowest, and reach a minimum in summer when temperature and irradiance are highest. (**B**) Zooplankton density peaks following spring and summer phytoplankton blooms. (**C**) Densities of target cyanobacterial taxa. (**D**) Cyanobacterial net growth rates from consecutive daily logged density measurements. (**E-F**) Convergent cross-mapping (CCM) analysis showing coupling strength between environmental variables and net growth rates of (**E**) *Microcystis* and (**F**) *Dolichospermum*. Orange bars represent coupling strength (convergence skill rho) between observed and predicted values from the original time series. Grey bars show mean coupling strength from 100 seasonal surrogate time series with red error bars indicating the standard error. Biotic variables used in **Figs. 3-5** are highlighted in bold. Strong seasonal coupling is marked by similar heights in orange and grey bars (e.g., for oxycline depth, temperature, irradiance, thermocline depth, NH_4_, and PO_4_). In contrast, strong daily-scale coupling is shown by orange bars exceeding grey bars.

### Cyanobacterial dynamics

Lake Greifensee is periodically affected by cyanobacterial blooms and associated cyanotoxins^18–20^ (**Fig. 1**). Our study focuses on two key groups: the *Microcystis* group (*Microcystis* sp., *Aphanothece* sp., *Aphanocapsa* sp.) and *Dolichospermum* sp. *Microcystis*, the most common potentially-toxic bloom-forming cyanobacterium worldwide^21^, typically blooms in Greifensee during summer and autumn (**Fig. 1C**). Previous studies confirm the production of microcystins and numerous other bioactive metabolites from this group in the lake, including during our study period^18,20^. *Dolichospermum*, the second most common potentially toxic bloom-forming cyanobacterium globally, is known to produce both hepato- and neurotoxins^3,22^. While rarely forming surface blooms in Greifensee, it reaches peak abundance in late summer, and bioactive metabolites potentially associated with this taxon are found during the study period^20^. Our high-resolution data reveal significant temporal fluctuations in both cyanobacteria (**Fig. 1C**). This provides a unique opportunity to examine growth dynamics and their relationships with biotic and abiotic factors at temporal scales directly relevant to bloom formation.

To quantify these dynamics and investigate the influence of biotic interactions, we present a growth rate analysis based on high-frequency monitoring data. To estimate daily growth rates, we calculate the difference in Ln-transformed densities for each taxon between consecutive days, as recorded by the underwater microscope (**Fig. 1D**). This approach reflects the net daily gains or losses observed for each target taxon, and it is conceptually similar to the per capita growth rate estimates obtained from laboratory studies^8,23^. For each taxon, nGR_i_=Ln(N)_i,t+1_ - Ln(N)_i,t_ where i=cyanobacterial taxon, N_t_=density at time t, N_t+1_=density at time t+1. Using tracked density observations from 2019 to 2023, we estimate the daily nGR for both *Microcystis* and *Dolichospermum*. While the nGR patterns explain the duration and intensity of cyanobacterial peaks in densities, we observe how the maxima in nGRs and the resulting overall densities varied considerably over the studied years (**Fig. 1, C and D**). These aperiodic fluctuations exemplify the hallmark characteristics of complex ecological systems like plankton communities: non-equilibrium and non-linear dynamics^24–26^.

The presence of these dynamics necessitates analytical approaches beyond conventional methods. Analysing such complex systems, where chaotic dynamics lead to bounded but aperiodic fluctuations with sensitive dependence on initial conditions, demands specialised modelling approaches. These methods must explicitly account for nonlinear and state-dependent relationships, including dynamic changes in growth rates^27–30^. For these reasons, we use state-space reconstruction methods, which do not rely on detailed mechanistic knowledge of the focal system and outperform traditional models in chaotic systems^31,32^, including plankton^26^, to study the high-frequency dynamics of cyanobacterial nGR as a function of their environmental conditions. We choose ecologically important environmental factors that are known to influence the physiology and interactions of planktonic organisms^33–35^, specifically cyanobacteria^3,6,36^. We demonstrate below how they relate to the nGRs of two globally important cyanobacteria taxa.

### Coupling to environmental factors

We first ask how strongly biotic and abiotic environmental factors are coupled to cyanobacterial nGRs in our field dataset. Plankton communities are forced by seasonal changes in temperature and nutrients (**Fig. 1, A to C**) that drive an ecological succession^34^. These abiotic factors, external to the plankton community, may create apparent synchrony (correlation) between non-interacting species that share similar environmental requirements^27^, or drive such interactions^17^. To estimate the strength of the direct coupling between abiotic/biotic factors and the nGR of cyanobacteria we use convergent cross mapping (CCM)^27^, with and without controlling for seasonal effects. Specifically, we compare the CCM rho (cross-mapping skill) value calculated using the original time series with 100 seasonal surrogate time series created by randomly sampling days from the original data across different years within the same calendar weeks (see **materials and methods**). Deterministic links at the daily scale that are significantly different from seasonal forcing are identified when at least 95% of the CCM predictive skill rho of the surrogate time series is smaller than that of the original time series.

Abiotic factors (such as NO_3_, NH_4_, PO_4_, water temperature and irradiance depth) measured in the upper water column (1-8 m) have a high CCM rho indicating a strong coupling with cyanobacterial nGRs. Still, their predictive skill is comparable to that of the seasonal surrogates, indicating no substantial difference between the two (**Fig. 1, E and F**). This suggests that their effects on cyanobacterial nGR represent seasonal forcing, as expected from the prominent seasonal fluctuations of the abiotic environment (**Fig. 1A**). The only abiotic variables that rank high in CCM rho after controlling for seasonal effects are the meteorological conditions (precipitation, wind speed, and air pressure), commonly used for modelling cyanobacterial blooms^10^ (**Fig. 1, E and F**). On the other hand, biotic factors, including the density of chrysophytes, diatoms, rotifers, calanoid copepods, cladocerans and overall cyanobacterial abundances, deviate substantially from the seasonal surrogates (**Fig. 1, E and F**). Biotic factors are strongly coupled at the daily scale with changes in cyanobacterial nGRs, a pattern we also observe in two more groups of cyanobacteria from Lake Greifensee, *Chroococcus* and *Coelosphaerium*, which are nontoxic (**Fig. S5**). This suggests that biotic factors are more important than abiotic environmental conditions in explaining bloom dynamics on a daily scale.

### Growth rate responses

After identifying the biotic factors that are more strongly coupled to cyanobacterial growth rates, we ask how the abundance of zooplankton and phytoplankton taxa shape nGR response patterns when interacting with seasonal drivers like PO_4_ and water temperature. To investigate this, we analyse the dependence of the nGR of *Microcystis* and *Dolichospermum* on the daily time series of 23 explanatory variables (environmental parameters and plankton densities) using the multiview distance regularised S-map method (MDR S-map)^28,37,38^. (**Figs. 1, A to D, and Fig. S4**). All explanatory variables were set at a one-day lag relative to the nGR to focus on the biologically relevant scale (day) at which growth (cell division) and loss (grazing, sinking) occur^39^. We use a multiview distance approach to address the high dimensionality^38^, elastic-net regularisation to mitigate collinearity and overfitting^37^, and test 125 S-map models with varying parameters (theta, alpha, lambda) to select the best fit based on sequential leave-future-out cross-validation (see **materials and methods**).

S-map models reveal strong nonlinearity (high theta parameter) as a defining feature in all six best models describing cyanobacterial nGRs (**Fig. 2, Fig. S6, and Table S4**). Our forecasting models outperform constant-rate baseline models over a 30-day horizon (**Fig. 2**) and maintain forecasting accuracy, even after controlling for temporal autocorrelation (**Figs. S7 and S8**). Prediction accuracy exceeds baseline performance from day 7 to day 15 (**Fig. 2**) in our target taxa and increases only by 2-4 days in the most conservative autocorrelation controls across all taxa (**Figs. S7 and S8**). This demonstrates that our models successfully capture temporal processes driving cyanobacterial nGR by incorporating key time-varying biotic and abiotic factors. The S-map coefficients for both biotic and abiotic factors show significant temporal variation, further confirming the dynamic nature of these interactions (**Fig. 3A and Figs. S9 to S12**). We consistently observe negative density dependence in cyanobacterial nGR, while environmental factors exhibit taxon-specific nonlinear effects (**Figs. S9 to S12**). These findings reveal that cyanobacterial bloom formation results from complex, dynamic interaction networks that shift across environmental gradients^17,23,38,40^—a critical insight for predicting blooms under climate change and eutrophication scenarios.

**Fig. 2:**
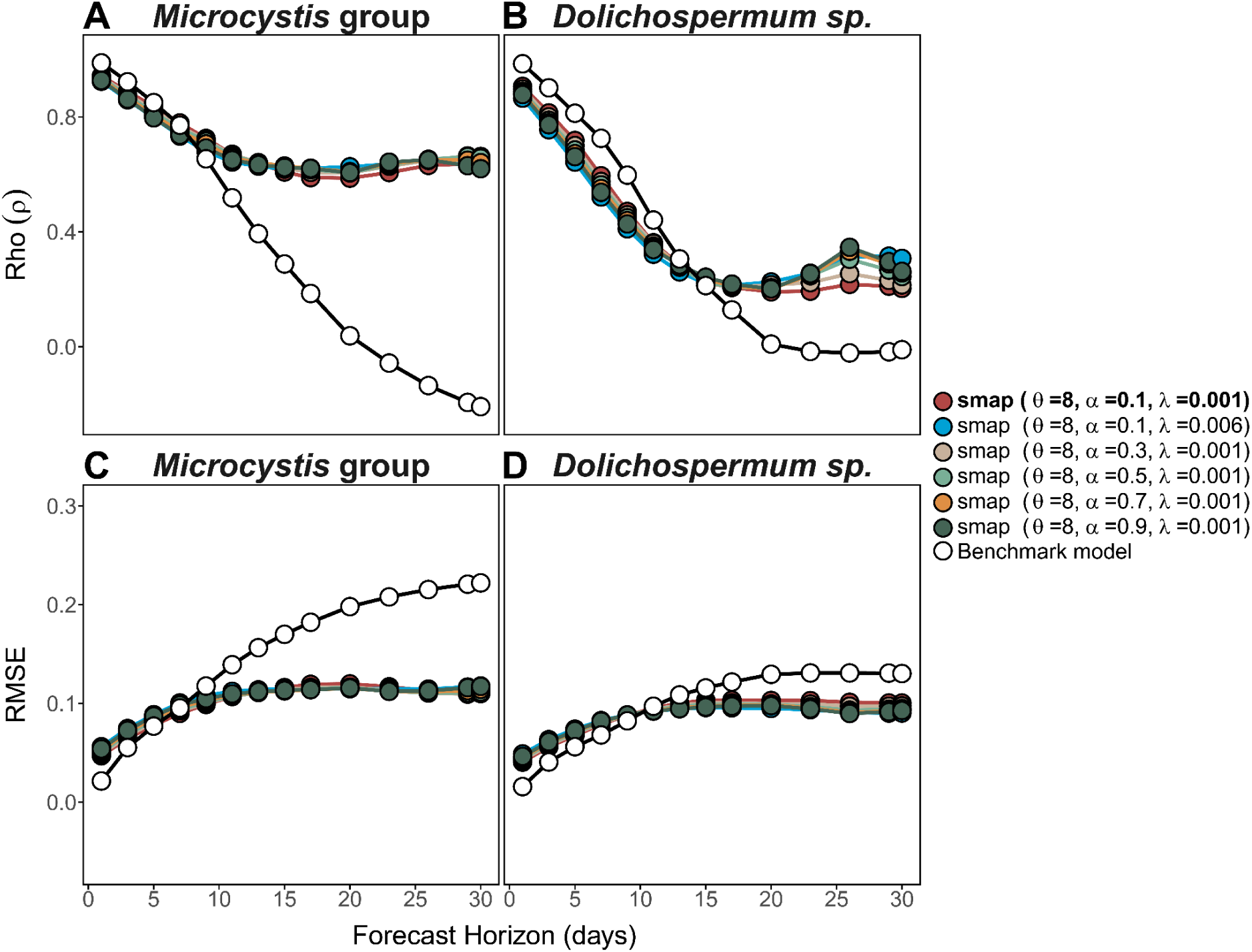
Forecast proficiency of MDR S-maps for *Microcystis* and *Dolichospermum* growth rates. Predictive performance of S-map models evaluated across increasing forecast horizons (1-30 days). The six best-performing parameter sets (selected via cross-validation at a 1-day horizon) are used for evaluation. Performance is quantified using (**A-B**) Pearson’s correlation coefficient rho (ρ) and (**C-D**) root mean squared error (RMSE) between observed and predicted values. Coloured lines represent individual S-map models with distinct parameter combinations; the best-performing model (used for interaction analysis) is highlighted in bold. The black line shows the performance of a benchmark persistence model (future values remain unchanged; see **materials and methods**). S-map models consistently outperform the persistence model across extended forecast horizons, even when temporal autocorrelation is excluded (**Figs. S7-S8**), indicating the successful capture of underlying system dynamics.

**Fig. 3:**
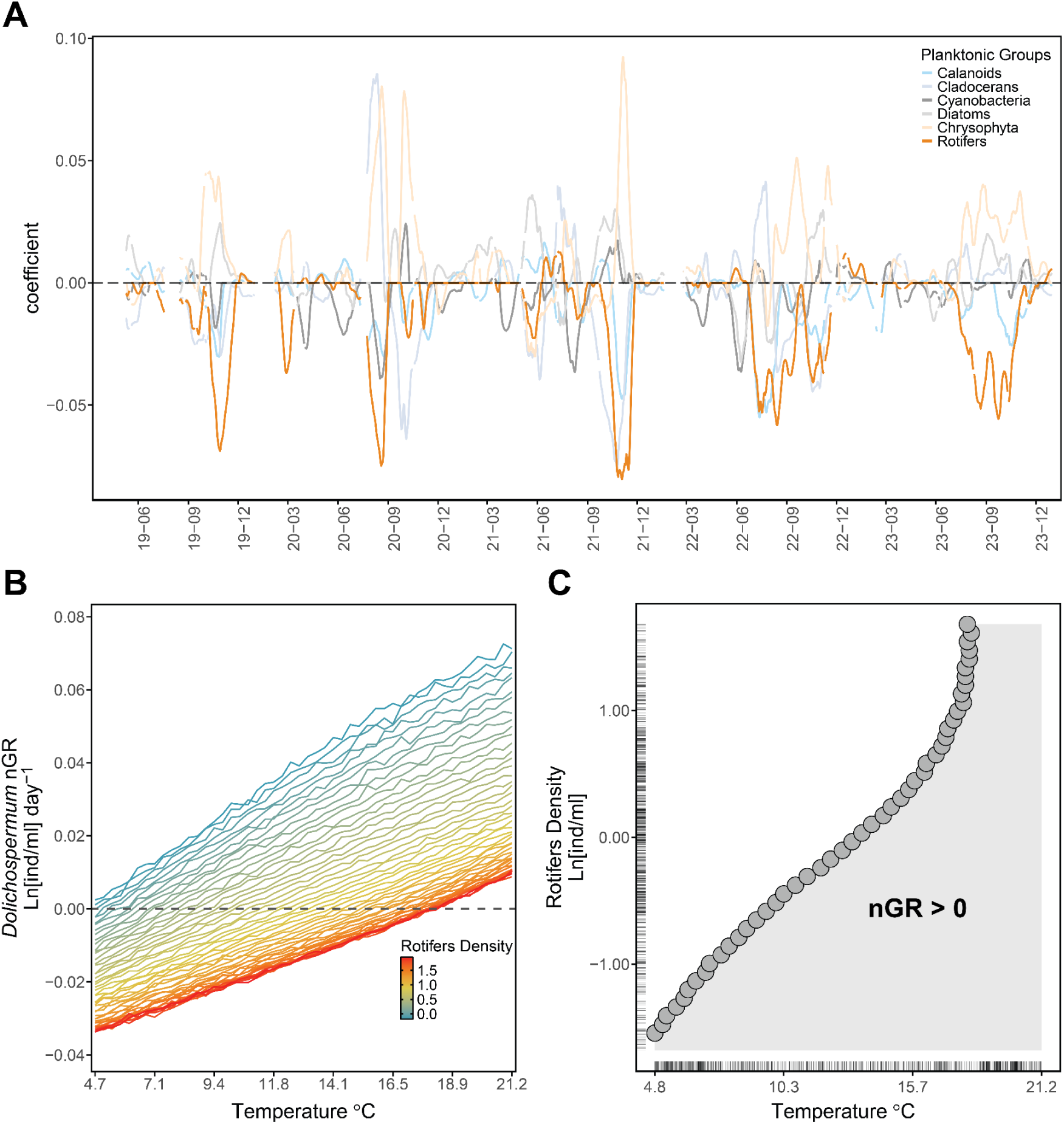
Time-varying biotic interactions and their effects on cyanobacterial blooming. (**A**) Temporal dynamics of pairwise interactions between *Dolichospermum sp.* and six key planktonic groups (calanoids, cladocerans, rotifers, cyanobacteria, diatoms, chrysophyta). Interaction coefficients are derived from MDR S-maps modelling of *Dolichospermum* net growth rate (nGR) (**materials and methods**). These biotic variables show high predictive skill (rho) in CCM analysis after seasonal effect removal (Fig. 1 **F**). Positive coefficients indicate growth facilitation, while negative coefficients suggest grazing pressure (e.g., rotifers—orange line) or competition. (**B**) Partial dependence analysis showing *Dolichospermum* nGR as a function of water temperature and rotifer scaled density. Color scale indicates rotifer density in Ln(ind/mL). The grey dotted line represents zero net growth (nGR=0); values above this line indicate a positive growth rate and biomass accumulation (blooming conditions). (**C**) Zero net growth analysis showing temperature-rotifer density combinations at which *Dolichospermum* transitions from negative to positive growth. The positive relationship demonstrates that higher rotifer densities require correspondingly high water temperatures for *Dolichospermum* to achieve blooming conditions. Grey surfaces indicate conditions supporting positive nGR (blooming). Tick marks on axes indicate the empirical density of temperature and rotifer observations in Lake Greifensee (2019-2023).

Having established the dynamic nature of these interactions, we next examine how specific biotic factors modulate cyanobacterial responses to key environmental factors. To unravel how biotic factors influence cyanobacterial growth over gradients of water temperature and dissolved inorganic phosphorus (PO_4_), we use MDR S-maps to predict the nGR of each focal cyanobacterium across observed conditions while holding other variables at their median values. In **Fig. 3B**, water temperature consistently promotes *Dolichospermum*’s nGR, while increasing rotifer density shifts the entire temperature response downward (as expected from a grazer) maintaining the same shape of the response curve but requiring higher temperatures for positive growth. We apply this approach systematically, examining how different biotic factors–like the abundance of zooplankton or other phytoplankton–modify cyanobacterial responses to both water temperature and PO_4_ gradients. While we observe the expected positive growth responses to increasing temperature and PO_4_ (**Figs. 4 and 5 and Figs. S13 and S14**), confirming their established role in bloom formation^3,5^, our analyses reveal how biotic interactions fundamentally alter these responses, effectively showing the environmental conditions that can trigger blooms.

**Fig. 4:**
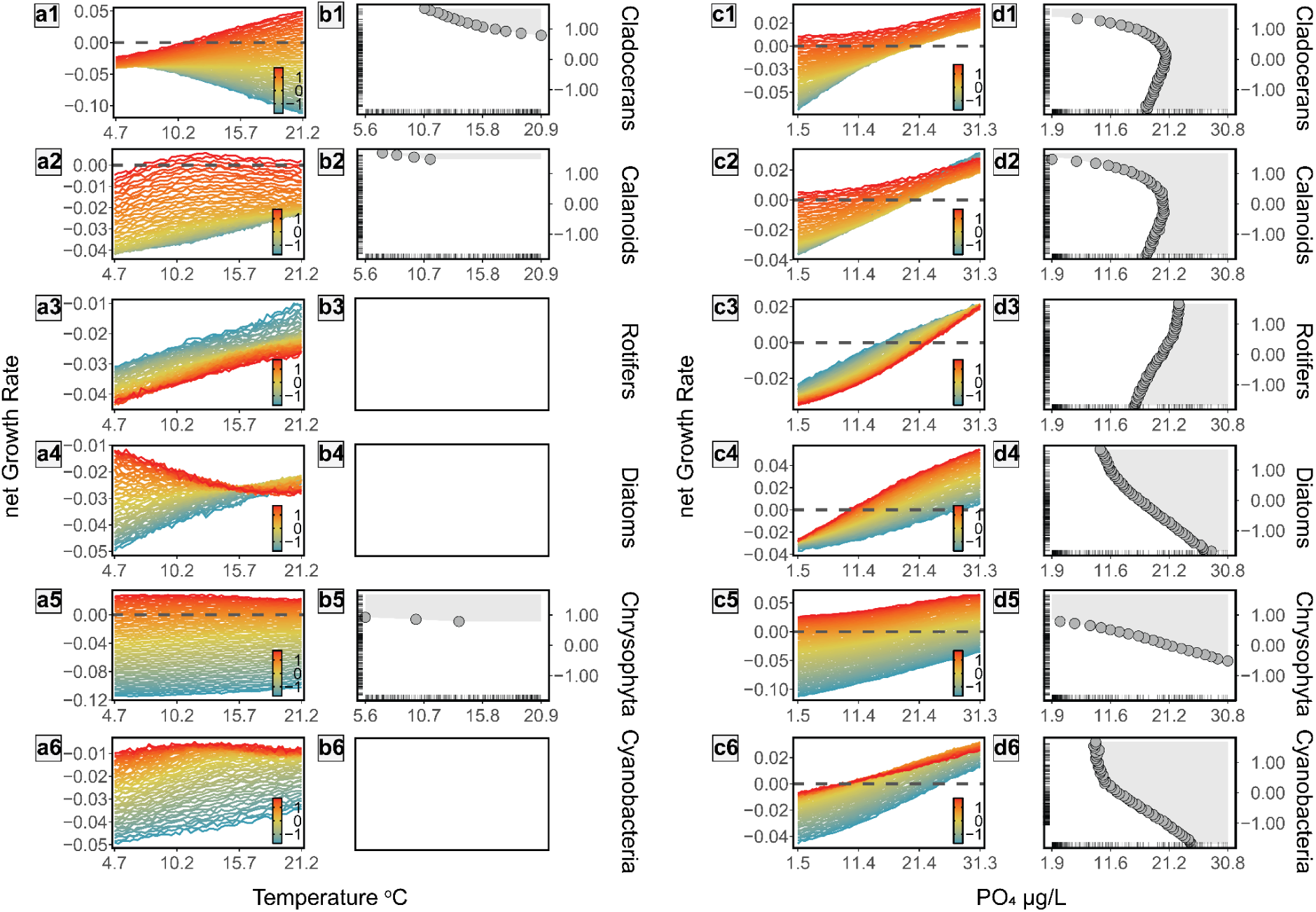
Biotic interactions shape *Microcystis* blooming across temperature and phosphate gradients. (**a1-a6**) Temperature dependency of *Microcystis* net growth rate (nGR, Ln(ind/mL)⋅day^−1^) across increasing densities (Ln(ind/mL)) of key planktonic groups: cladocerans (**1**), calanoids (**2**), rotifers (**3**), diatoms (**4**), chrysophytes (**5**), and other cyanobacteria (**6**). (**c1-c6**) Phosphate (PO_4_) dependency of *Microcystis* nGR across densities of the same planktonic groups. (**b1-b6, d1-d6**) Zero net growth analysis showing blooming conditions along temperature and PO_4_ gradients as influenced by increasing densities of interacting organisms. Grey surfaces indicate conditions supporting positive nGR (blooming). Tick marks on axes show empirical distributions of temperature, PO_4_, and planktonic group scaled densities in Lake Greifensee (2019-2023). The relationship between blooming thresholds and organism density reveals either bloom inhibition (positive correlation indicates higher temperature/PO_4_ requirements for blooming as organism density increases) or bloom facilitation (negative correlation indicates lower temperature/PO_4_ requirements as organism density increases). White boxes (**b3**, **b4**, **b6**) indicate cases where zero net growth analysis was not possible because *Microcystis* had negative nGR across all tested conditions (see Y scale on panels **a3**, **a4**, **a6**). Detailed pairwise interaction analyses in **Figs. S15** and **S19**.

**Fig. 5:**
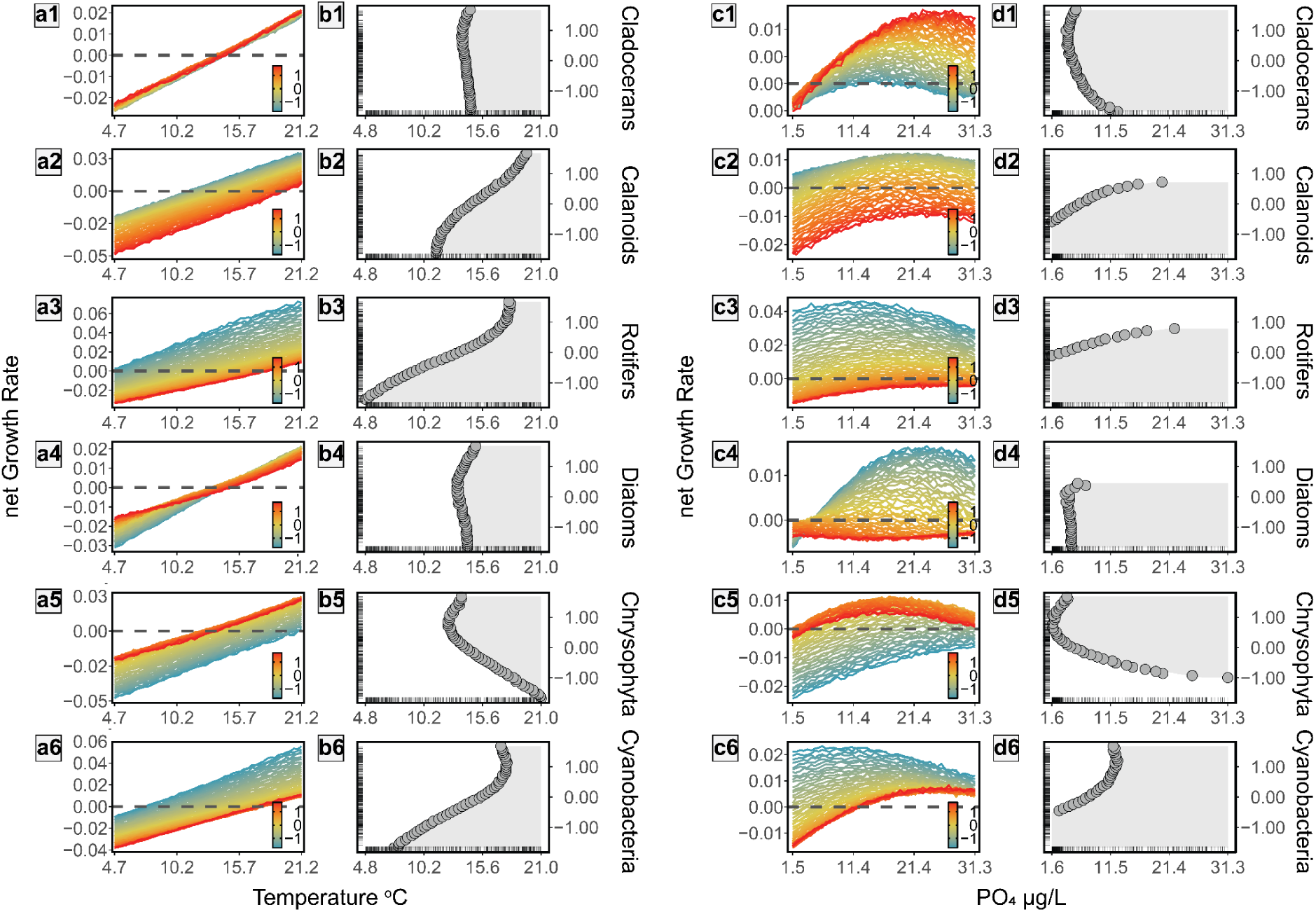
Biotic interactions shape *Dolichospermum* blooming across temperature and phosphate gradients. (**a1-a6**) Temperature dependency of *Dolichospermum* net growth rate (nGR, Ln(ind/mL)⋅day^−1^) across increasing densities (Ln(ind/mL)) of key planktonic groups: cladocerans (**1**), calanoids (**2**), rotifers (**3**), diatoms (**4**), chrysophytes (**5**), and other cyanobacteria (**6**). (**c1-c6**) Phosphate (PO_4_) dependency of *Dolichospermum* nGR across densities of the same planktonic groups. (**b1-b6, d1-d6**) Zero net growth analysis showing blooming conditions along temperature and PO_4_ gradients as influenced by increasing densities of interacting organisms. Grey surfaces indicate conditions supporting positive nGR (blooming). Tick marks on axes show empirical distributions of temperature, PO_4_, and planktonic group scaled densities in Lake Greifensee (2019-2023). See Fig. 4 for a detailed interpretation of the relationships between blooming thresholds and organism densities. Detailed pairwise interaction analyses in **Figs. S16** and **S20**.

Building on our analytical approach of mapping growth response patterns across environmental gradients, we further show that focal biotic factors (those deviating significantly from seasonal surrogates - **Figs. 1, E and F, and Fig. S5, C and D**) can fundamentally alter both the magnitude and shape of cyanobacterial growth responses to temperature and PO_4_ (**Figs. 4 and 5 and Figs. S13 and S14**). Remarkably, we observe cases where established positive growth responses to temperature and PO_4_ completely reverse direction under specific biotic conditions. For instance, increasing cladoceran abundance shifts *Microcystis’* temperature response from negative to positive, while increasing diatom abundance causes the opposite shift (**Fig. 4, a1 and a4**). Similar biotic effects on growth response patterns occur across PO_4_ gradients (**Figs. 4 and 5**) and all other measured abiotic conditions (NH_4_, NO_3_, oxycline depth, air pressure–**Figs. S15 to S18**). While laboratory studies previously demonstrated changes in phytoplankton response curves with varying abiotic conditions^41,42^, our study provides the first field evidence of biotic factors reshaping these fundamental growth responses. These findings suggest that ignoring dynamic biotic interactions fundamentally mischaracterises bloom formation, potentially leading to errors in predicting both the timing and magnitude of cyanobacterial blooms.

### Cyanobacterial realised niche

To assess how plankton taxa influence cyanobacteria’s ability to achieve net positive growth (their realised niche^43,44^), we identify the temperature and PO_4_ thresholds where cyanobacteria cross the zero net growth isocline (nGR = 0) and begin accumulating biomass (**Fig. 3B**). We then plot these zero nGR thresholds against the abundance of biotic factors (**Fig. 3C**). For example, **Fig. 3C** shows how increasing rotifer abundance (a potential grazer) elevates the temperature requirements for *Dolichospermum* blooming. Our results reveal that both zooplankton and phytoplankton abundances significantly alter the temperature and PO_4_ requirements for positive growth in both focal cyanobacteria (**Figs. 4 and 5 and Figs. S13 and S14**). The magnitude of these biotic effects is substantial—shifting blooming requirements by several °C and µg/L of PO_4_—especially considering Lake Greifensee’s mean annual temperature of 12.9°C (SD = 5.8°C) and mean PO_4_ of 9.0 μg/L (SD = 11.2 μg/L) in the upper water column. We find consistent evidence that biotic interactions shape the realised niche of *Microcystis* and *Dolichospermum* (**Figs. 4 and 5**), as well as *Chroococcus* and *Coelosphaerium* (**Figs. S13 and S14**). Importantly, the effects of biotic factors on cyanobacterial realised niches vary in a taxon-specific manner across temperature and PO_4_ gradients (**Figs. 4 and 5 and Figs. S13 and S14**). These findings highlight how species-specific biotic interactions fundamentally structure cyanobacterial ecology, suggesting that models that neglect these interactions will likely misrepresent population dynamics and fail to accurately predict the environmental conditions favouring blooms.

*Microcystis*, a cyanobacterium thriving in eutrophic environments and warm, stable waters^6^, is primarily favoured by increasing temperature and phosphorus levels^11,45,46^. Our field observations confirm these abiotic drivers’ positive effects on *Microcystis* accumulation (**Fig. 4 and Figs. S15 and S19**). Beyond these abiotic factors, our analysis reveals complex biotic interactions significantly influencing *Microcystis* growth rates. Most zooplankton groups—including calanoids, cyclopoids, cladocerans, ciliates, and copepod nauplii—enhanced *Microcystis* nGR across water temperature and PO_4_ gradients, effectively shifting response curves upward and expanding the realised niche (**Fig. 4 and Fig. S15**). This facilitation likely occurs through indirect mechanisms, such as selective predation on competing species^13^. Rotifers, however, emerged as a notable exception, consistently suppressing *Microcystis* nGR across all temperature and PO_4_ levels, suggesting direct grazing pressure^15^.

Phytoplankton interactions display temperature-dependent patterns. Diatoms and chlorophytes exhibit a shift from facilitation at low temperatures to competition at high temperatures, while chrysophytes show particularly strong facilitation effects, enabling *Microcystis* growth at PO_4_ concentrations as low as 3.3 μg/L (**Fig. 4, c5 and d5**). Quantitatively, high densities of facilitating groups (chrysophytes, calanoids, cladocerans, cyanobacteria, and diatoms) reduced the PO_4_ requirements for *Microcystis* blooming on average by 19.68 μg/L. In contrast, high rotifer densities increased PO_4_ requirements from 15.56 to 22.9 μg/L, further confirming their inhibitory effect (**Fig. 4 d3**) (see **materials and methods**). Notably, facilitation—one of the least studied biotic interactions in bloom dynamics—appears critically important for *Microcystis* response to PO_4_. We interpret these facilitation mechanisms through multiple pathways: directly via mutualistic interactions between cyanobacteria and photosynthetic protists^47^, and indirectly either by diverting grazers toward more palatable organisms (e.g., diatoms and chlorophytes)^13^ or through selective mixotrophic suppression of competing protists (e.g. by chrysophytes)^48^. These findings highlight how positive biotic interactions may substantially expand *Microcystis* bloom potential beyond the limitations predicted by abiotic factors alone.

In contrast to *Microcystis*, *Dolichospermum* prefers moderate nutrient levels and shows more variable responses to environmental conditions in Greifensee^6^. Our analysis reveals that biotic interactions predominantly constrain *Dolichospermum*’s realised niche, shifting response curves downwards and inhibiting blooming (**Figs. 5 and Figs. S16 and S20**)—a marked difference from the generally facilitative interactions observed with *Microcystis*. Most plankton groups exhibit inhibitory effects on *Dolichospermum* accumulation. Rotifers demonstrate the most dramatic impact, substantially increasing environmental thresholds for blooming by raising temperature requirements from 4.78 to 18.22°C and PO_4_ requirements from 1.64 to 22.3 μg/L (see **materials and methods**). Copepods similarly exhibit strong negative effects.

Despite this predominantly inhibitory pattern, two groups emerge as notable facilitators: cladocerans and, particularly, chrysophytes. Consistent with their effects on *Microcystis*, chrysophytes show remarkable positive effects on *Dolichospermum*, substantially lowering blooming thresholds. In their presence, temperature requirements decrease from 21 to 12.41°C, while PO_4_ requirements drop from 31.28 to 1.79 μg/L (**Fig. 5, b5 and d5**). This facilitation extends beyond *Dolichospermum* and *Microcystis*, as similar patterns were observed for *Chroococcus* (**Fig. S13**). The strong facilitation by chrysophytes appears particularly crucial for *Dolichospermum*, as their abundance could determine whether environmental conditions support blooming events (**Fig. 5**). These findings highlight chrysophytes as potential key taxa in cyanobacterial bloom dynamics, suggesting that monitoring them may significantly improve bloom forecasting models beyond traditional abiotic predictors.

### Implications

Understanding how biotic interactions shape species’ realised niches remains a fundamental challenge in ecology^49^. Our work provides an empirical demonstration of how realised niches emerge from species interactions—a fundamental concept applicable across ecological systems from microbes to animals and forests with significant implications for both ecological theory and management practice. Biotic interactions in plankton communities can dramatically alter blooming requirements across multiple environmental dimensions: facilitative relationships enable blooms at unexpectedly low nutrient levels, explaining occasional bloom occurrences under seemingly unfavourable conditions, while temperature thresholds can shift by 10-13°C—potentially advancing or extending seasonal bloom windows by months in temperate lakes. Conversely, antagonistic interactions through grazing or competition can prevent blooms entirely, even when abiotic conditions appear favourable. This evidence resolves longstanding paradoxes in bloom prediction failures. Critically, we identified specific planktonic groups with consistent effects across cyanobacterial taxa—particularly rotifers, copepods, and cladocerans as natural bloom regulators, and chrysophytes as potential bloom facilitators—suggesting that monitoring programs should incorporate these plankton groups as early warning indicators of cyanobacterial blooms, and that targeted food web manipulations^50,51^ could complement traditional nutrient management approaches for more effective bloom control strategies. While our empirical data come from Lake Greifensee, the fundamental mechanisms we identify—biotic modification of environmental thresholds—represent a universal ecological process applicable to freshwater systems globally.

### Limitations

Our study reveals key biotic interactions shaping cyanobacterial net growth, though some aspects of microbial ecology remain unexplored. While we capture major planktonic interactions, the roles of parasites and heterotrophic bacteria on nGRs warrant further investigation. For instance, the cyanobacterial microbiome likely influences nutrient availability and growth dynamics^52^, and viruses or fungi may significantly impact loss processes^51^. As expected in complex ecological systems, our findings demonstrate context dependency in cyanobacterial responses. While specific interaction patterns may vary across lakes and under future conditions—particularly with novel perturbations like species invasions—the mechanistic understanding of how biotic factors modify cyanobacterial niches represents fundamental knowledge. The empirical approach we developed to quantify biotic effects on bloom thresholds provides a broadly applicable framework for monitoring and managing cyanobacterial blooms across diverse freshwater ecosystems, even though particular species interactions differ.

### Conclusions

Climate change and eutrophication are driving increasingly frequent and severe cyanobacterial blooms, demanding improved prediction and management. Our study reveals that biotic interactions fundamentally shape bloom formation conditions—a critical insight missing from current conceptual models and forecasting approaches. Current models of cyanobacterial blooms are built on a fundamental assumption: they treat environmental thresholds as fixed properties of species when our evidence demonstrates these thresholds are emergent properties of complex biotic interactions. Collecting and analysing high-frequency data of cyanobacteria and co-occurring plankton species, we demonstrate that these interactions significantly modify bloom requirements across environmental gradients. Neglecting these biotic effects in predictive models risks underestimating bloom expansion under climate change, particularly for *Microcystis*, whose niche seems to expand through facilitative mechanisms. Our findings offer practical advances: incorporating high-frequency plankton data into monitoring programs can provide early-warning signals that complement traditional water quality measurements. Additionally, targeted manipulation of key planktonic groups—such as rotifers or chrysophytes—presents novel management strategies beyond nutrient control. By integrating these biotic dimensions into forecasting models, monitoring and management approaches, water resource managers gain valuable tools to anticipate and mitigate the substantial economic and ecological costs of harmful cyanobacterial blooms.

## Materials and Methods

### 1. Data acquisition

#### 1.1 Study System (Lake Greifensee)

We used high-frequency data collected from a monitoring platform in Lake Greifensee, Switzerland, from March 2019 to December 2023. Lake Greifensee is a mesotrophic lake with a surface area of 8.45 km², a maximum depth of 32 metres, and an average depth of 18 metres. The lake’s elevation is 435 metres above sea level and has a volume of 0.15 km³. Due to its relatively deep profile and temperate climate, Lake Greifensee exhibits a monomictic mixing regime, with complete water turnover occurring annually in winter. The lake is classified as eutrophic (with an average annual level of 0.04 mg/L of total P), and has a long history of nutrient pollution and toxic cyanobacterial blooms^18–20^.

#### 1.2 Measured Parameters

The tracked variables included the lake’s physical, meteorological, and chemical parameters. Plankton community data are described in Section 2.4. The physical parameters of the lake –water temperature, photosynthetically active radiation (PAR, wavelengths 400-700 nm) and dissolved O_2_– were measured using an Ocean Seven CTD probe (https://www.idronaut.it/) over the water column at 0.1 m intervals (from 1 to 17 m) with a sampling frequency of 3 to 6 (wintertime) hours. Weather conditions, such as maximum wind speed, precipitation and air pressure data, came from a weather station installed on the lake monitoring platform (Vaisala Oyj WXT520 & OTT HydroMet WS700-UMB) every 10 min. To assess nutrient concentrations (P-PO_4_, N-NO_3_, and NH_4_), we collected water samples at the monitoring station from a depth of 3 metres once per week and analysed them with standard methods for nutrient chemistry^53^.

### 2. Data Processing

#### 2.1 Water Physics

Water temperature data were restricted to the photic zone (0-8 m), above the thermocline (**Fig. S4**), where plankton thrive and image acquisition occurs. Due to potential shading from the monitoring platform, we used PAR light penetration depth as a proxy for light intensity. This was the depth at which PAR reached the detection limit (5 μEm^−^²s^−1^)^54^. Oxycline depth was defined as the depth at which dissolved oxygen reached 4 mg/L. Thermocline depth was calculated using the R package rLakeAnalyzer (v1.11.4.1). All CTD data were averaged to obtain daily values.

#### 2.2 Lake Meteorological Conditions

We averaged data of maximum wind speed, precipitation and air pressure from the weather station installed on the lake monitoring platform to daily estimates.

#### 2.3 Water Chemistry

Measurements of the inorganic nutrients nitrate (NO_3_), ammonium (NH_4_) and phosphate (P-PO_4_) were performed weekly with water samples collected at 3 m depth. To obtain nutrient levels at a daily frequency, we predicted daily nutrient values using a Random Forest (RF) model trained and validated for each nutrient^54^. Nutrient levels fluctuate daily mostly due to variations in water’s physical parameters and meteorological conditions^35^. Interpolation of nutrient values based on autocorrelation of the time series would not account for such a daily variation. We, therefore, trained RF models on weekly nutrient data (n = 201) based on CTD and Meteo information from the same date, tested and validated these nutrient models on unseen nutrient values (10-times cross-validation where 80% of the data were used for training and 20% for testing), and then used RF models to predict (as opposed to interpolate) daily nutrient concentrations based on daily data of water physics and meteorological conditions (**Table S5**). Random forest model training, validation and testing were performed using the Caret package in R (v6.0-94).

#### 2.4 Plankton Data

##### 2.4.1 Automated Image Analysis & Image Classification

Plankton was monitored with a dual-magnification dark-field imaging microscope that captures all particles in the size range between ∼10 μm and ∼ 1 cm at 3 m depth using two magnification objectives: one for phytoplankton and small zooplankton (5.0x) and one for larger zooplankton (0.5x)^55,56^. The instrument is currently manufactured by www.guatek.com. The choice of deployment depth in Greifensee is motivated by the average structure of the lake water column during the growing season: the thermocline is generally around 8 m (**Fig. S4**) and the epilimnion is generally well mixed by daily thermal winds. The 3 m depth is generally a representative sample of the phytoplankton community of the photic zone^54^. Images were collected at a rate of 1 frame/s for 10 minutes every hour and regions of interest (ROIs) were automatically identified by the instrument using a Canny edge detector^56^. A Python image processing script performed colour conversion, edge-detection, and morphological feature extraction (e.g. object dimensions) of detected raw ROIs (https://github.com/tooploox/SPCConvert). These objects were manually annotated and classified by 2 trained taxonomists (authors MR and SE), and served as input for the image classifier, after being resized to 224 x 224 pixels (Imagenet standardisation)^57,58^.

A hierarchical classification structure was employed, consisting of two classifiers for the 5.0x magnification: the former distinguishes between different cyanobacteria species and a background class, while the latter further classifies samples predicted to be of the background class into additional phyto- and microzooplankton classes. This improves the accuracy of cyanobacterial classification by reducing the false positive predictions. A separate classifier was trained to distinguish images in the 0.5x magnification. All three classifiers shared the same pre-trained BeITv2 architecture^59^ but differed in the last layer. The classifiers were trained in two stages, in the first stage only the weights of the input and output layers of the neural network are optimised. In the second stage, all weights are unfrozen and optimised. The AdamW optimiser with a weight decay of 0.01 was used, and the learning rate was scheduled with linear warmup cosine annealing^60^. A validation set (15%) was used for checkpoint selection, and the training set (70%) was augmented to improve generalisation, the remaining 15% are used as a test set on which metrics given in the following are computed (**Figs. S1 to S3 and Table S3**). The first classifier was trained using a Focal Loss with γ=2 and equal weight for both classes^61^ to obtain high precision on cyanobacteria classes, while Categorical Cross Entropy was used for the other two classifiers^61^. Test-time augmentation was used during inference to improve robustness^59^. The performance of the three classifiers can be seen in **Figs. S1 to S3**. For further details on the image classification visit https://github.com/kaechb/lit_ecology_classifier.

##### 2.4.2 Transformation of Planktonic Densities

Classified object counts at the daily scale were Log-transformed (base 10) and smoothed using a local polynomial regression to remove random noise as described in Section 2.5 (loess function with a span of 0.03). For aggregation of taxa into higher taxonomic groups across different magnifications and to facilitate comparison, Log10-transformed ROI/s values were converted to Log10 individuals/mL using calibration functions described in^56^ (for the 5p0x magnification, Log(ind/mL) = -2.9 + 0.71 Log(ROI/s); for the 0p5x magnification, Log(ind/ml) = -0.91 + 1.3 Log(ROI/s)). Following unit transformation, densities were back-transformed (anti-Logarithm) and taxa were aggregated within and/or across magnifications to create higher taxonomic groups for our analysis (see below). Finally, these groups were transformed with the natural Logarithm for further analysis (e.g. estimation of growth rates). A total of 1742 data points representing taxon and group-specific density estimations were collected over five years (after removing days when the underwater microscope was serviced or there were not enough images), spanning March 2019 to December 2023.

##### 2.4.3 Target Taxa & Planktonic Groups

Our target cyanobacteria included two potentially toxin-producing taxa, the *Microcystis* group (including *Microcystis* sp.*, Aphanothece* sp.*, Aphanocapsa* sp.) and *Dolichospermum* sp., and two non-toxic taxa, *Chroococcus* sp. and *Coelosphaerium* sp. To assess the influence of biotic interactions on cyanobacteria, we aggregated zooplankton and phytoplankton taxa into broad taxonomic groups. From the 34 classified zooplankton taxa, we grouped taxa according to their phylogenetic class into 6 categories (**Table S1**), capturing distinct zooplankton feeding behaviours. Similarly, the 49 identified phytoplankton taxa were aggregated to the phylum level, resulting in 6 taxonomic groups (**Table S2**).

#### 2.5 Smoothing of daily data

Daily data of all variables used in this study were smoothed using a local polynomial regression (loess in Stats, v3.6.2) function with a span of 0.03 to remove serially uncorrelated (random) noise. The chosen span was verified by examining the autocorrelation in the residuals after smoothing to ensure no removal of deterministic signals from the time series: **Figs. S21 and S22** show the distribution of residuals after smoothing, **Figs. S23 and S24** their partial autocorrelation function.

### 3. Data Analysis

All analyses and figure production were conducted in R version 4.4.1.

#### 3.1 Convergent Cross-Mapping

To assess the strength of coupling between environmental variables and cyanobacterial net growth rate (nGR), we tested for direct links between them using convergent cross-mapping (CCM)^27^. We applied CCM to test the strength of causal links between environmental variables and cyanobacterial nGR, estimating the forecast skill (rho–Pearson’s correlation coefficient between observations and predictions–Section 3.5) at Tp=0 (prediction horizon) to focus on direct interactions. To factor out potential seasonal influences on the strength of causal links, we conducted CCM on 100 seasonal surrogates. These surrogates were created by randomly substituting values within each calendar week of the target year with values from the same week in other years (4 or 5 years depending on the chosen month). Finally, we calculated the forecast skill by extracting the seasonally estimated linkages computed from the surrogate time series and compared these values to those obtained from the original time series, across all tested links. The three zooplankton and phytoplankton groups that were the strongest coupled to the target cyanobacterial nGRs were selected for further analyses (**Fig. 1, E and F and Fig. S5, C and D**). The CCM function was used with an embedding dimension of 15, a prediction horizon of 0, an exclusion radius of 16 and 80% of the original library size (rEDM, v1.15.4).

#### 3.2 nGR modelling with S-map

We modelled the nGR of the 4 target cyanobacterial taxa using the multiview distance regularised S-map method (MDR S-map) ^17,28,37,38^. S-maps belong to the EDM framework and were chosen because of their capacity to infer interactions in highly non-linear, high-dimensional and noisy data describing natural plankton communities. S-maps are locally-linear multiple regression models, where data points are assigned varying weights based on their position in state-space. State-space dependence allows interactions between variables to change over time as the system changes states, avoiding the assumption of stationary interactions. The interactions between the target and explanatory variables are estimated through the coefficients of the S-maps model at each time point. All explanatory variables were scaled (with the scale function in R) before fitting regularised MDR S-maps (rEDM, v0.4.7).

#### 3.3 Dealing with High Dimensionality

For modelling, we used 23 explanatory variables (or embedding dimensions), including 10 ecologically relevant physical, chemical, and meteorological parameters and 13 biotic variables representing the densities of observed planktonic groups that could potentially interact with cyanobacteria (6 zooplankton, 6 phytoplankton groups and the density of each target taxon). To mitigate the challenges posed by high dimensionality and avoid inaccuracies in nearest-neighbour distance calculations within state-space, we employed a multiview approach^28,38^ to generate weighted average distance estimates from the top 100 out of 1000 multiview embeddings. Each multiview embedding was constructed by randomly selecting 9 variables from the available 23. To evaluate these 1000 multiview embeddings, we utilised simplex projections with leave-one-out cross-validation and an exclusion radius of 14 points. The 100 configurations that yielded the lowest root mean squared error (RMSE) in forecast skill were selected and the highest-performing multiview embeddings were assigned greater weight in the distance calculation^28^. Distances between points in state-space were computed using Euclidean distance with the dist function in R (vegan, v2.6-8). In addition, to address potential issues arising from the use of numerous explanatory variables (collinearity) and noise in the data, which could compromise our estimates of interaction coefficients and lead to overfitting, we implemented elastic-net regularisation^37^. This technique penalises the model coefficients and improves the generalisation of the model (model selection). Elastic net regularisation was applied using the glmnet function in R (glmnet, v4.1-8).

#### 3.4 Parameter Selection and Model Validation

The regularised multiview distance S-maps require the a priori selection of 3 important tuning parameters, theta, alpha and lambda. Theta is a parameter within S-maps that controls the weights assigned to data points during local linear regression fitting. Alpha and lambda are parameters of the elastic net regularisation function. Alpha regulates the balance between the L1 (Lasso) and L2 (Ridge) penalties (α=1 corresponds to Lasso, and α=0 corresponds to Ridge) while lambda controls the overall strength of the penalty.

To select the optimal combination of these tuning parameters, we tested all possible combinations of the following values: theta (0, 0.1, 1, 3, 8), alpha (0.1, 0.3, 0.5, 0.7, 0.9), and lambda (1.000, 0.178, 0.032, 0.006, 0.001), resulting in 125 models (**Fig. S6**). The predictive performance of each model was evaluated with a sequential leave-future-out cross-validation, whereby starting with an initial library size of 10%, we predicted one time point into the future and progressively added the predicted point to the training set, continuing this process until the end of the time series. The best model parameters were the ones with the lowest root mean squared error (RMSE) in forecast skill one day forward (**Table S4**).

#### 3.5 Forecasting proficiency of nGR models

To evaluate the predictive power of our best-performing model, used to study nGR responses, we compared its forecasting proficiency with the next five top models (a total of six models with different combinations of theta, alpha, and lambda parameters, as described in Section 3.4). In this forecasting exercise, each model (of the four target cyanobacterial taxa) was used to predict the future nGR across forecast horizons (*TP*) ranging from 1 to 30 days using the sequential leave-future-out approach (as described in Section 3.4). At the same time, we assessed the models’ sensitivity to autocorrelation in the time series by systematically excluding an increasing number of the closest time points when generating predictions (exclusion radius–*ER*). This sensitivity analysis is particularly relevant for short-term forecasts. We set *ER* = *TP + n*, where *n* ranged from 0 to 15 days: in this way *ER* ≥*TP* to prevent data leakage into the model fitting. The forecast proficiency of each model was assessed using RMSE and Pearson’s correlation coefficient rho:

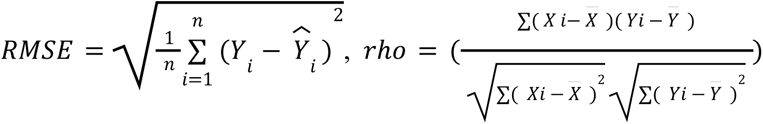

Forecast proficiency was evaluated relative to a persistence model, which assumes that the current growth rate of each cyanobacterium remains constant over the forecast horizon. In **Fig. 2** we present the forecasting results for the six best-performing S-map models with *n* = 0 (*ER = TP*), while **Figs. S7 and S8** display the mean and standard deviation of RMSE and ρ of the best six models across the full range of *n* values from 0 to 15.

#### 3.6 Main and Interactive Effects of Variables on Cyanobacterial nGR

To estimate the main effects of all explanatory variables, we extracted the coefficients from the MDR S-maps model fitted to the entire dataset with the optimal parameters. This resulted in 23 time series of coefficients (**Figs. S9 to S12**), which describe how the effect of one variable changes as the system evolves and other variables vary accordingly. To estimate the interactive effects of biotic and abiotic variables on cyanobacterial growth (multivariate partial dependence), we predicted nGRs allowing only the two interacting variables of interest (one biotic and one abiotic) to vary, while the rest were maintained at their median levels (**Figs. S15 to S18**). For each variable of interest, we selected a grid of 50 values spanning from the minimum to the maximum observed in the dataset, excluding the 10% of extreme values^17^. This resulted in 250 combinations of biotic and abiotic values for each multivariate partial dependency. To ensure the robustness of our response function patterns, we conducted a sensitivity analysis on the length of the data used to train the model. For this analysis, we extracted the response functions for each of the 250 combinations of values by randomly subsampling 50 times different proportions of the initial data (10%, 20%, 40%, 60%, 70%, 90%, and 95%). This additional step confirmed that the observed patterns of response functions were consistent across varying data lengths, enhancing the robustness and generalisability of our findings (selected examples for temperature and PO_4_ **Figs. S25 to S32**). In **Figs. 3C, 4 and 5 and Figs. S13 to S18**, we report the average of the 50 resamplings using 70% of the dataset for each of the 250 combinations of biotic and abiotic values.

#### 3.7 Quantifying the Effect of Biotic Interactions on Cyanobacterial Blooming

To quantify the effects that biotic factors have on the cyanobacterial zero net-growth isoclines (i.e. the blooming point), we focused on the conditions under which each target cyanobacterium achieves a net positive nGR. We predicted the multivariate partial dependencies of cyanobacterial nGRs (Section 3.6) as a function of the most common and strongly coupled biotic factors (Section 3.1), and two abiotic variables, water temperature and PO_4_, which are strongly but seasonally coupled to the cyanobacterial nGR and are widely recognised as important drivers of cyanobacterial blooms. The blooming point (intercept of the zero net growth isocline, nGR=0) is the value of water temperature or PO_4_ concentration above which cyanobacteria can accumulate biomass under a specific density of the target biotic group (**Fig. 3**). An increase in temperature or phosphorus levels that is required to attain a positive nGR with increasing biotic density suggests inhibition of blooming (**Fig. 3, B and C**). This could be interpreted as grazing pressure from zooplankton or competition by phytoplankton. Conversely, a decrease in the required temperature or phosphorus level required for growth with increasing density of biotic factors suggests a facilitation of blooming, regardless of the biotic group (**Fig. 4, c4 and d4**). To compare the effect of biotic factors within and between cyanobacteria, we calculated the range of variation in the blooming point (zero net growth isocline), as the difference between its maximum and minimum estimated values over temperature and phosphorus gradients for the target biotic factor (**Table S6**). For the effects of all biotic factors over all the abiotic factors on the blooming point of cyanobacteria, see **Figs. S19, S20, S33 and S34**.

#### 3.8 Data Availability

The raw data supporting this study’s findings are openly available from the Eawag Research Data Institutional Repository (ERIC/Open). Raw abiotic data can be found at https://doi.org/10.25678/000C8P and raw biotic data can be found at https://doi.org/10.25678/000C2G.

#### 3.9 Code Availability

The R scripts used to generate the results of this manuscript will be made available through the Eawag Research Data Institutional Repository (ERIC/Open) https://opendata.eawag.ch/, together with the final aggregated dataset.

## Acknowledgements

We thank Rudolf Rohr and Elisabeth Janssen for their valuable input and critical feedback. *Funding:* This research was funded by the Swiss National Science Foundation (project 202290 Cyanobloom), with the contribution for infrastructure by the Swiss Federal Office for the Environment (contract Nr Q392-1149).

## Author contributions

P.N. and F.P. conceptualised the study, and all authors contributed to its design. S.E., E.M., and P.N. prepared the data processing pipeline and assembled the data. P.N. performed the analyses. S.E., M.R., S.D., and M. B.-J. managed and annotated images, B.K. trained the machine-learning classifier and classified the images. P.N. and F.P. drafted the manuscript; all authors provided critical and conceptual feedback (contributing roles based on https://casrai.org/credit/).

## Supplementary Information

Figs. S1 to S34

Tables S1 to S6

