## Supplementary Information for "Biotic interactions shape the realised niche of toxic cyanobacteria"

**The PDF file includes:**

Figs. S1 to S34
Tables S1 to S6

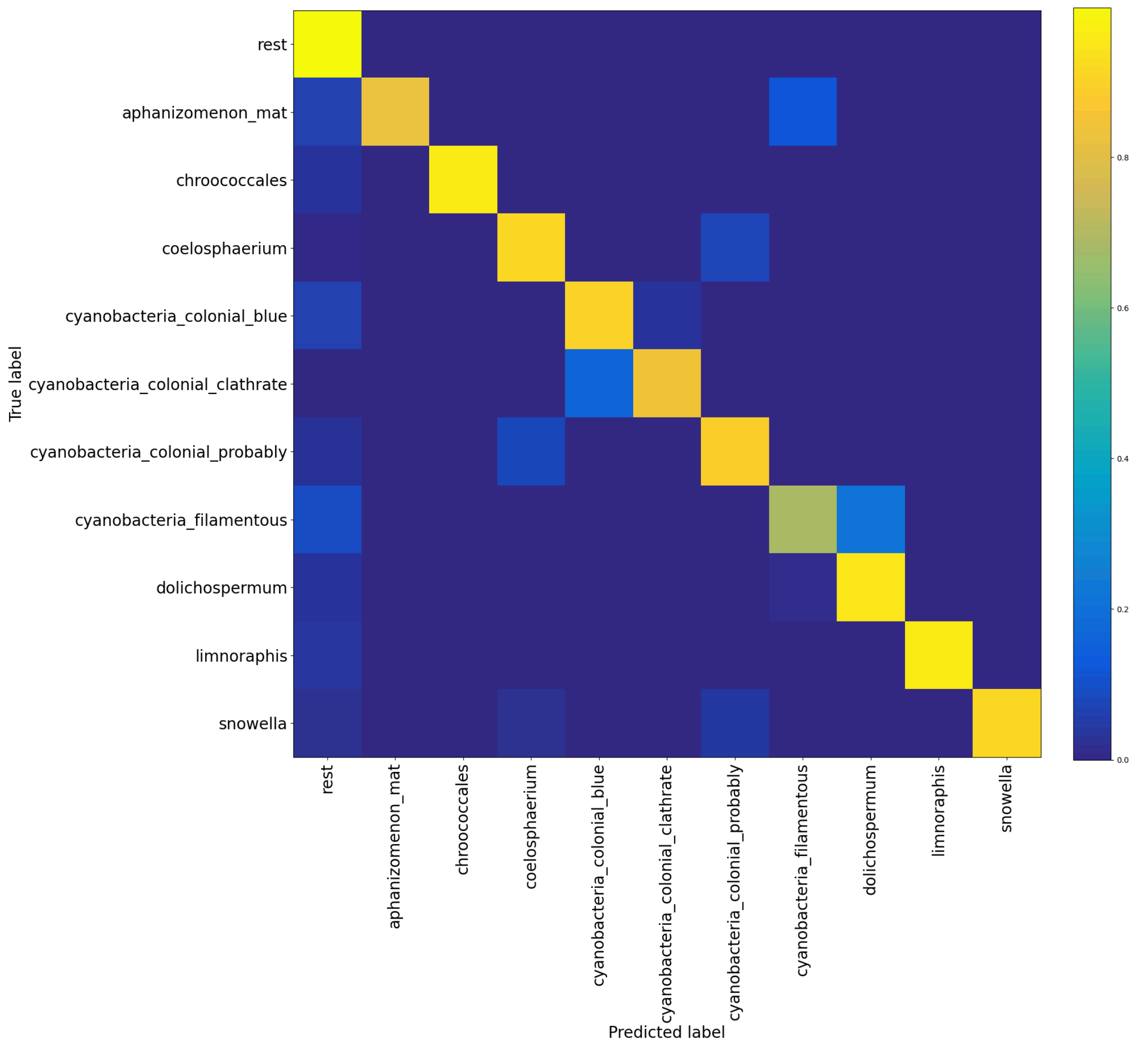


**Fig. S1: Confusion matrix of the Cyanobacteria Classifier.** Confusion matrix illustrating the performance of the cyanobacteria classifier designed to categorise 10 cyanobacteria classes from the rest in the 5.0x camera images. Training size: 27000 images; Validation size: 5785; Test size: 5787.

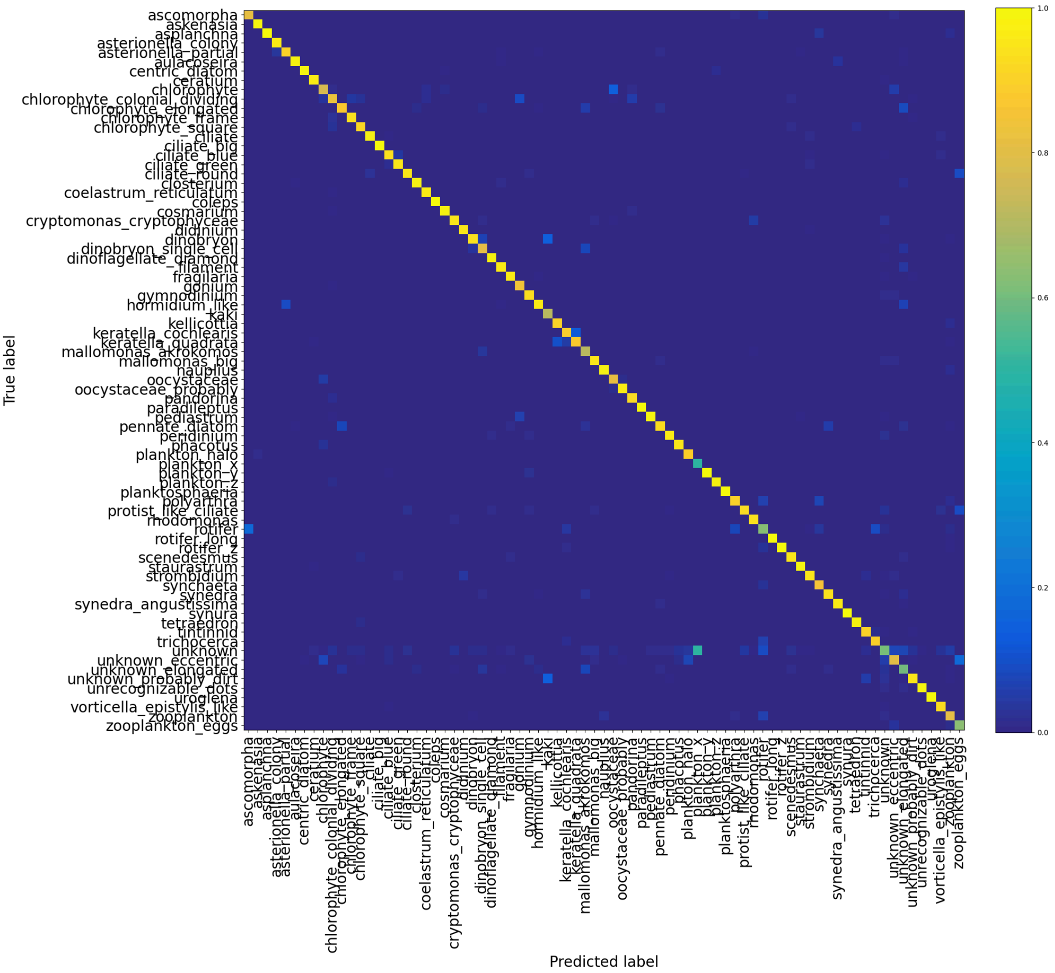


**Fig. S2: Confusion matrix of the Phytoplankton Classifier.** Confusion matrix illustrating the performance of the phytoplankton classifier designed to categorise the remaining classes in the 5.0x camera images following the implementation of the first Classifier. Training size: 24298 images; Validation size: 5206; Test size: 5208.

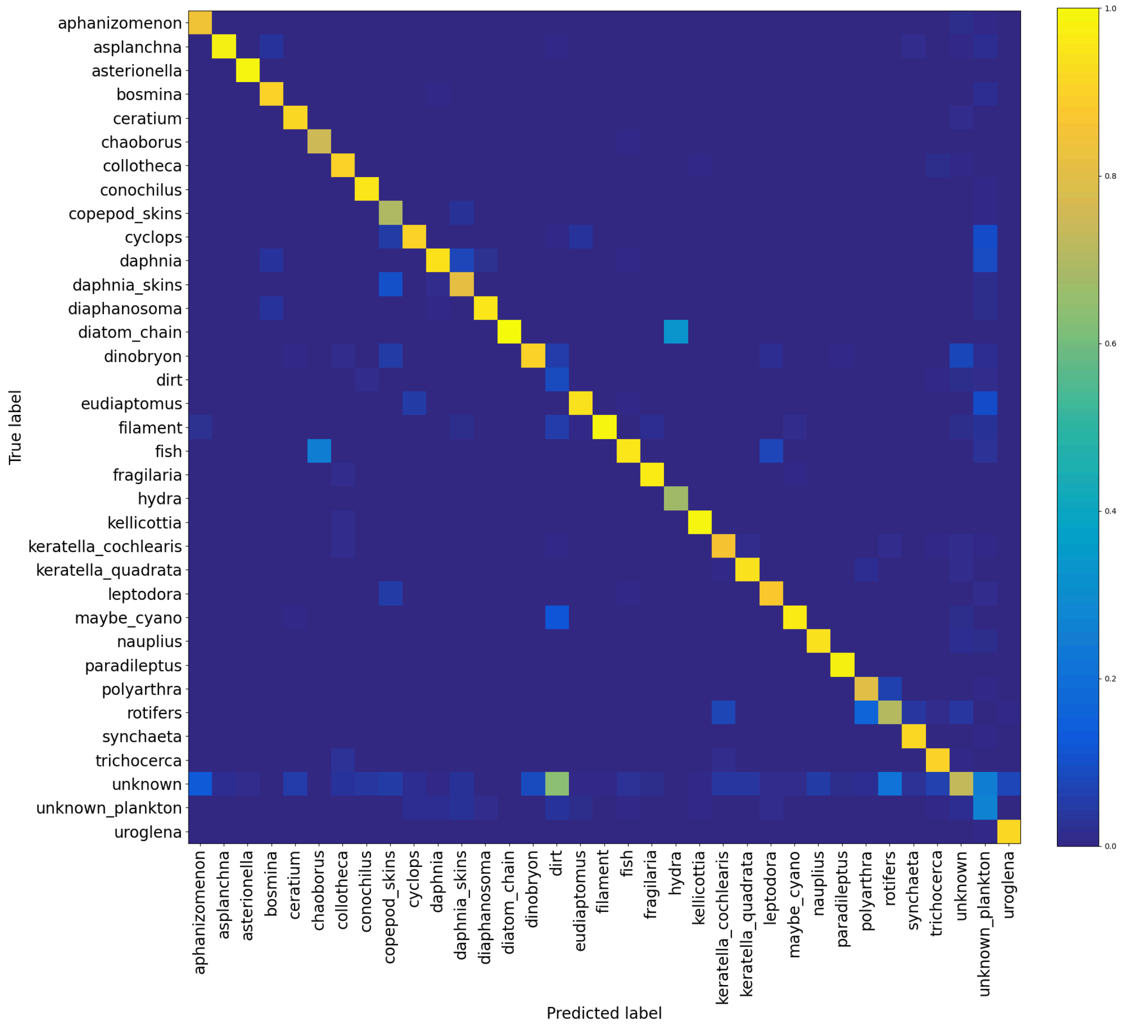


**Fig. S3: Confusion matrix of the Zooplankton Classifier** Confusion matrix illustrating the performance of the zooplankton classifier designed to categorise images of the 0.5x camera. Train size: 22680 images; Validation size: 4860; Test size: 4861.

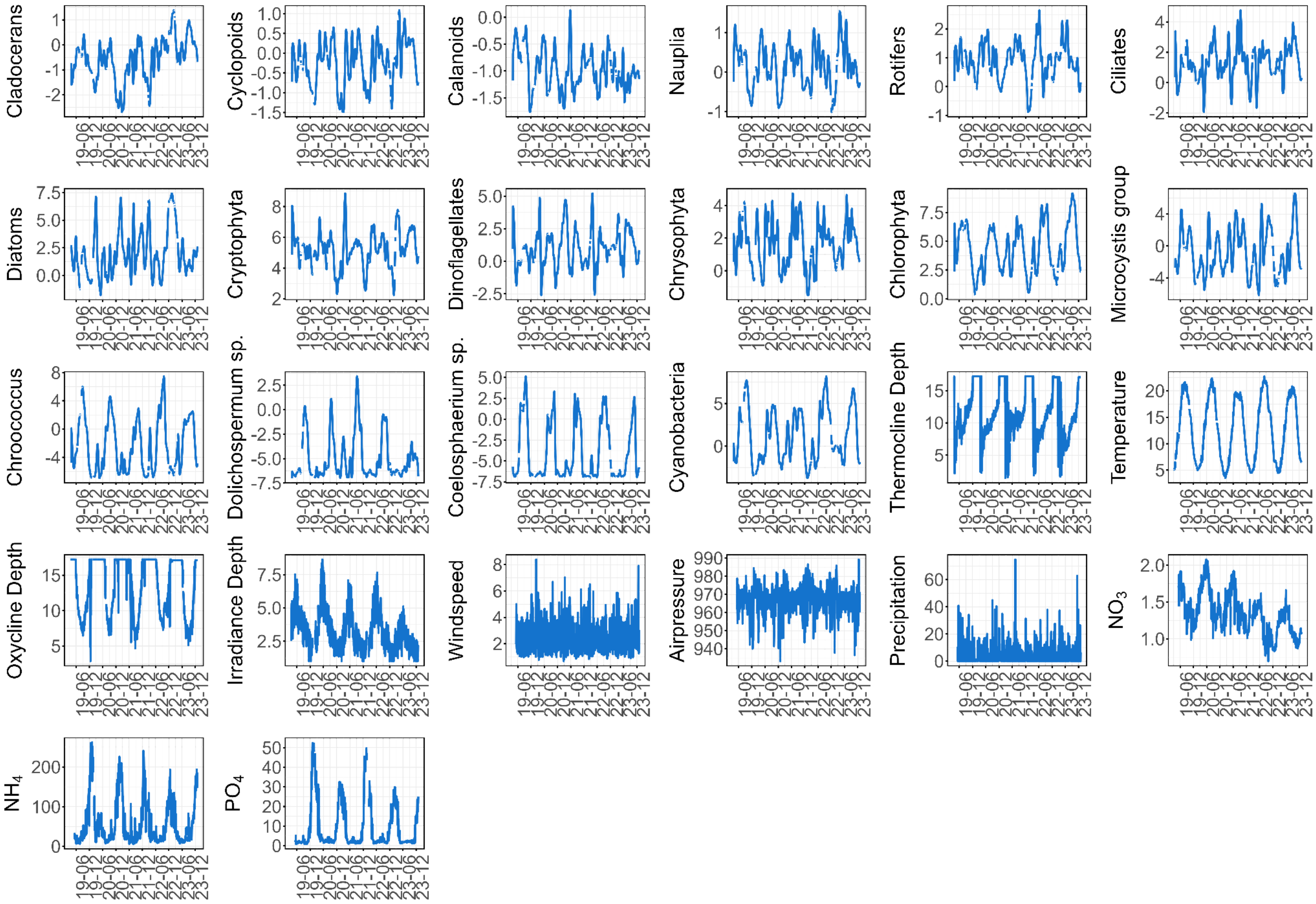


**Fig. S4: Time series of biotic and abiotic predictors used in our study.** Time series are smoothed with a loess function (span=0.03). Units as in **Fig.S15**.

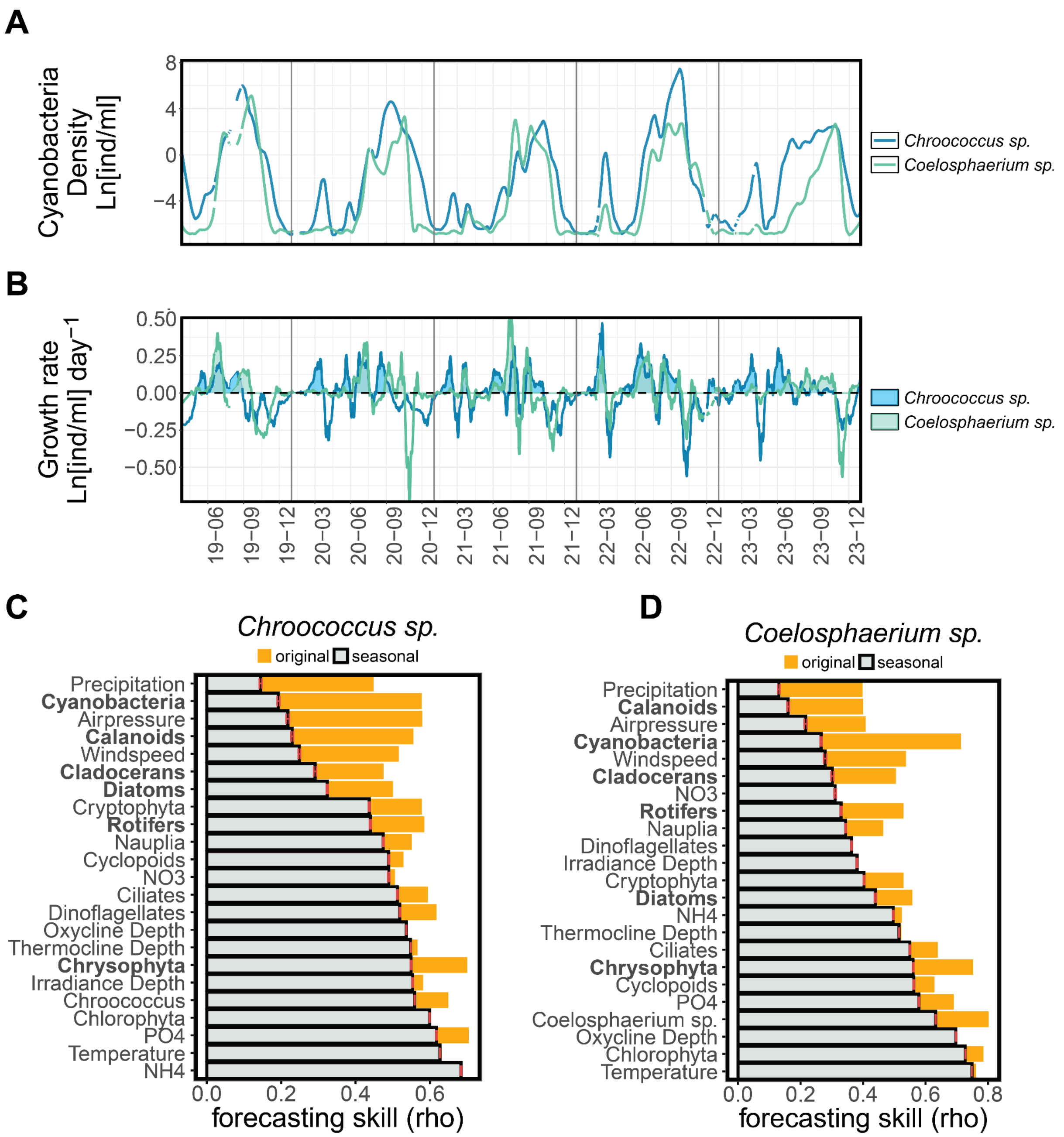


**Fig. S5: Temporal Dynamics and Biotic Coupling of *Chroococcus* sp. and *Coelosphaerium* sp. in Lake Greifensee.** (**A**) Target cyanobacteria densities; (**B**) net growth rates calculated as the difference in logged density between two consecutive days. Convergent cross-mapping (CCM) analysis between environmental variables (biotic and abiotic) and nGR of (**C**) *Chroococcus* sp. and (**D**) *Coelosphaerium* sp. Orange bars indicate coupling strength (convergence skill rho) between observed and predicted values from the original time series. Grey bars show the mean strength using 100 seasonal surrogate time series and red error bars show the SE estimate of the mean (see **materials and methods**). Biotic variables used for interaction analyses are highlighted in bold. High CCM rho values for oxycline depth, temperature, irradiance, thermocline depth, $NH_4$, and $PO_4$, combined with similar heights of orange and grey bars, indicate strong seasonal coupling rather than daily-scale interactions for these variables.

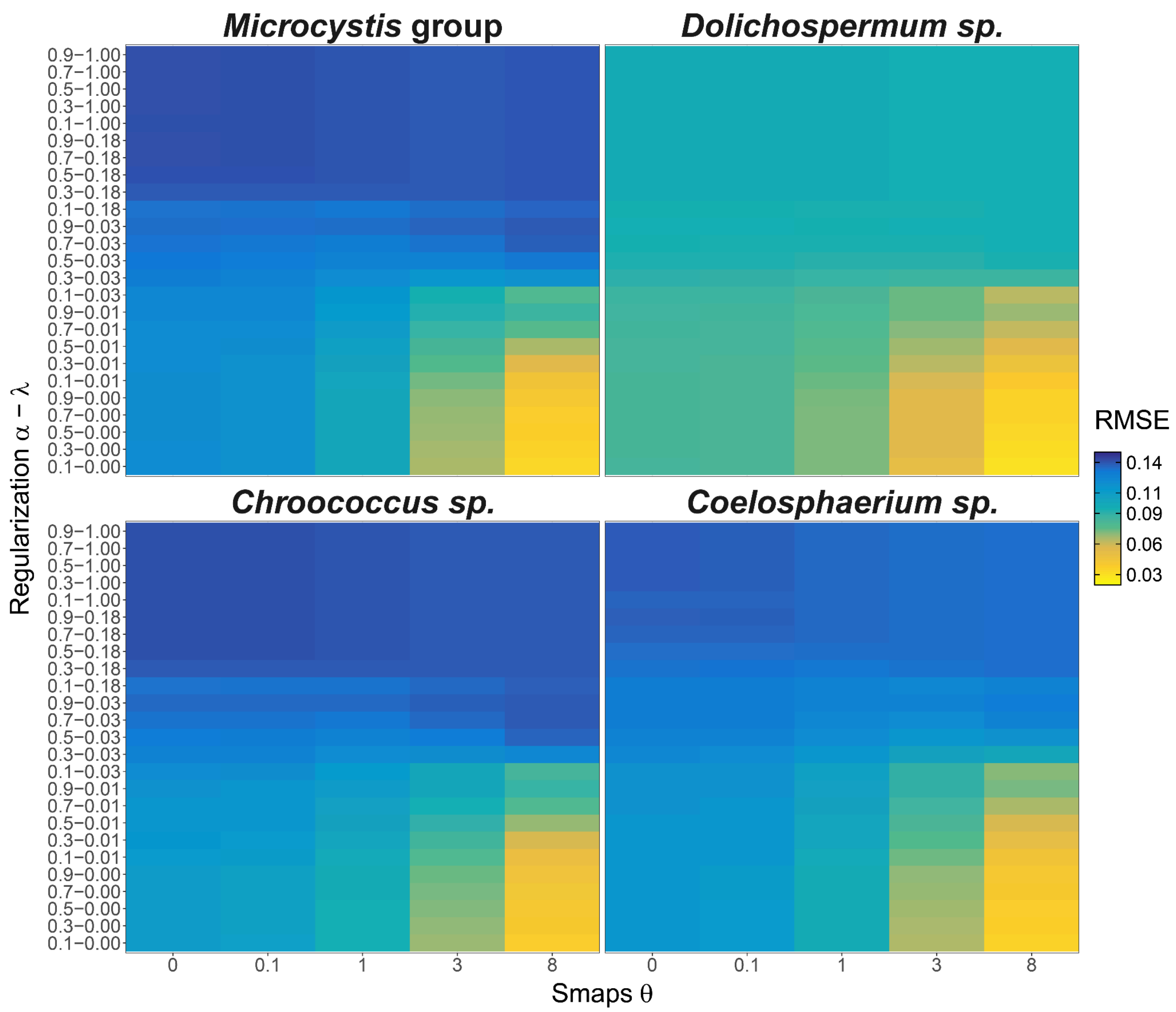


**Fig. S6: Performance of MDR S-maps for all parameter combinations tested in the models (θ, α, λ) for all target cyanobacteria nGRs.** The results indicate that MDR S-map models with θ = 8 achieve the highest predictive skill, measured as root mean squared error (RMSE). Colors indicate the predictive skill (RMSE) obtained from leave-future-out cross-validation, with warmer colors (e.g., yellow) representing higher predictive accuracy and cooler colors (e.g., blue) indicating lower accuracy. See **materials and methods** for a detailed explanation of the model parameters and the implementation of the cross-validation analysis.

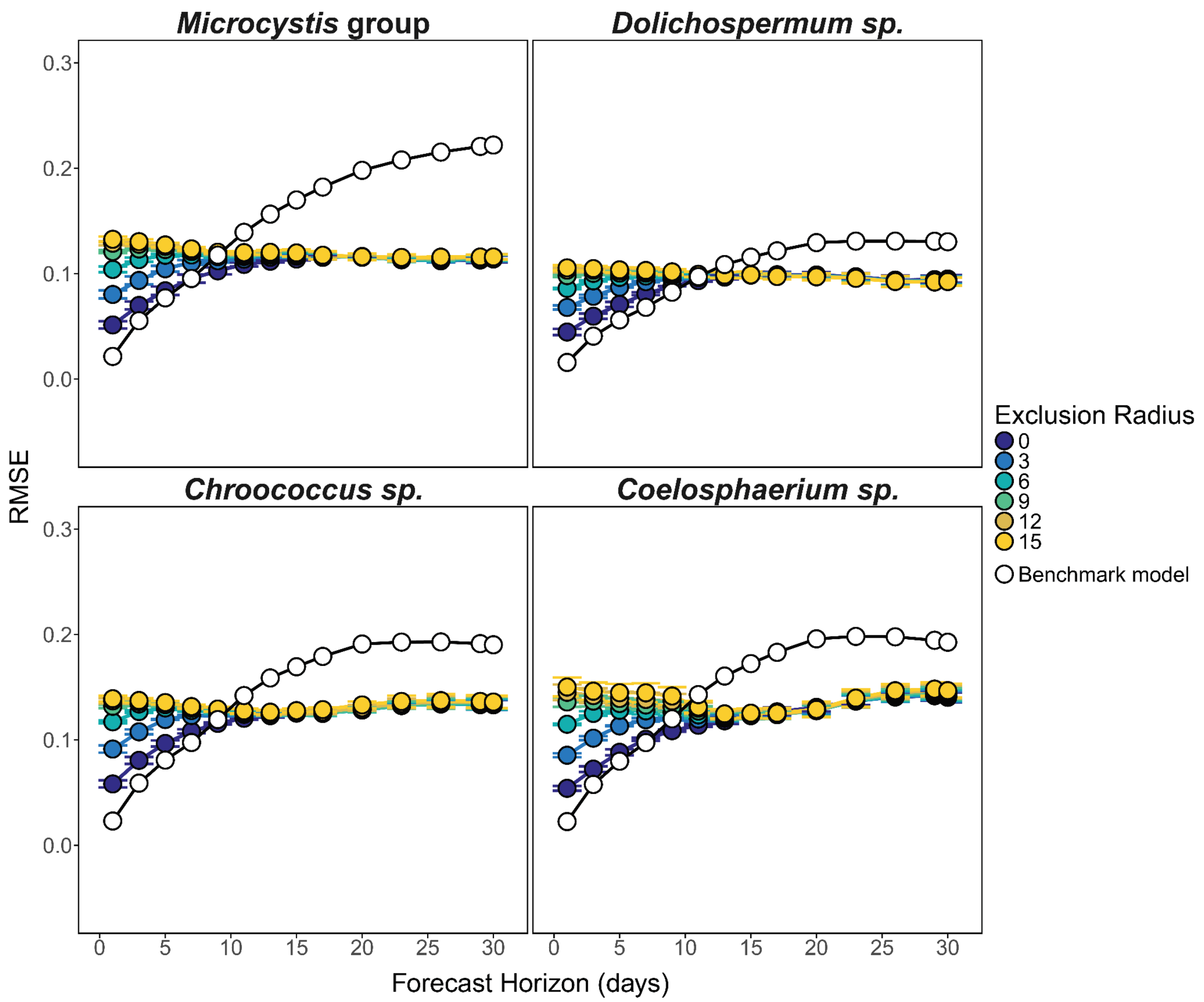


**Fig. S7: Forecasting error of MDR S-maps for cyanobacterial growth rates.** The predictive accuracy of S-map models was assessed across increasing forecast horizons (TP) and exclusion radii (ER), where ER was defined as TP+n (n={0,15}) to account for autocorrelation effects, for each target cyanobacterial group. The six best-performing parameter sets (selected via sequential leave-future-out cross-validation at a 1-day horizon) were used for evaluation. Model performance was quantified as the root mean squared error (RMSE) between observed and predicted values across forecast horizons from 1 to 30 days. Coloured points and lines represent the mean RMSE of the best-performing S-map models and error bars show the standard deviation. The black line represents the performance of a benchmark persistence model, which assumes that future values remain unchanged (see **materials and methods** for details). Notably, the S-map models successfully capture the underlying system dynamics beyond autocorrelation effects, as their predictive performance remains stable even at a 30-day forecast horizon. This robustness is further supported by the predictive capacity indicated by performance metrics such as Pearson's correlation coefficient, rho (see **Fig. S8**).

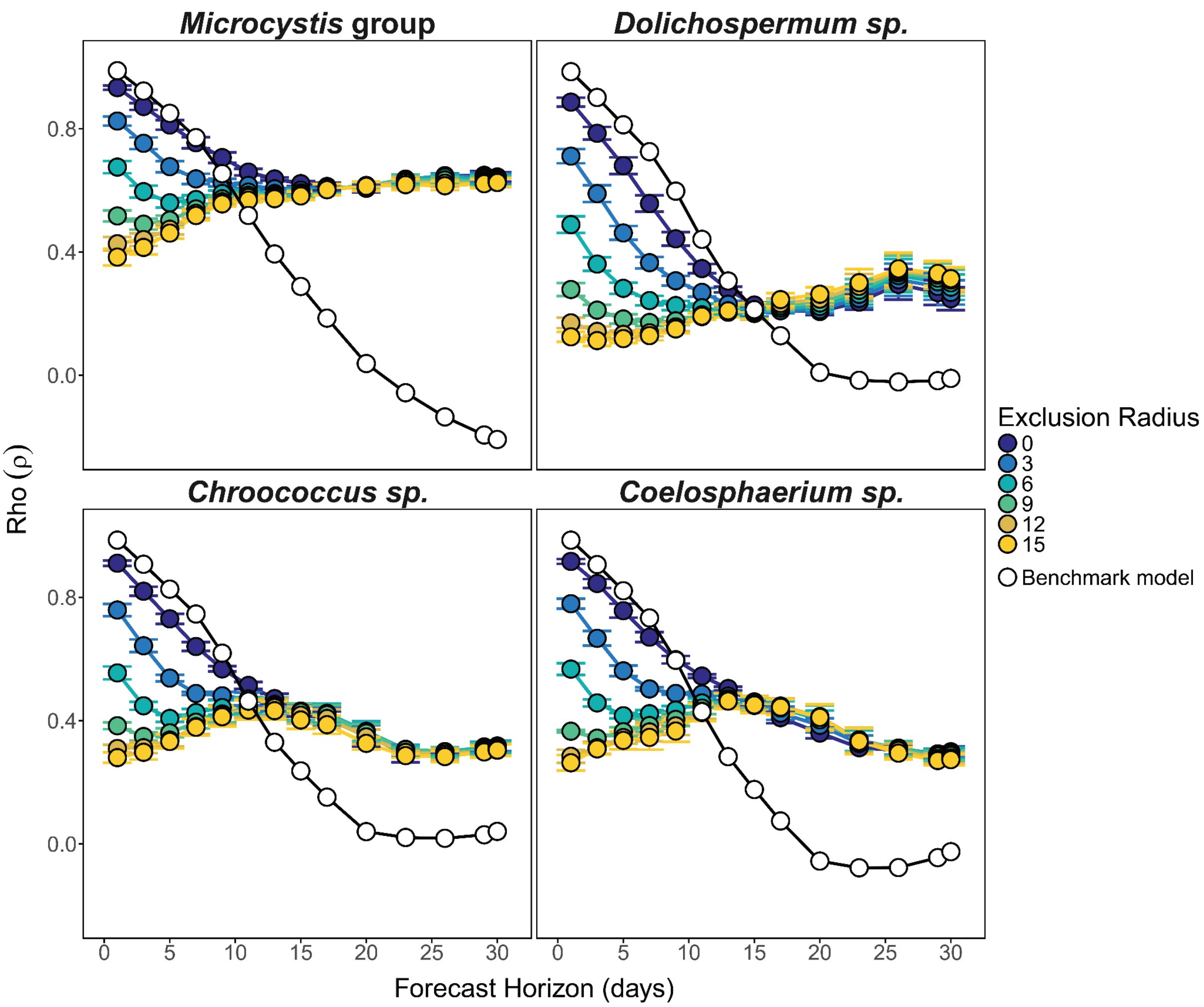


**Fig. S8: Forecasting skill of MDR S-maps for cyanobacterial growth rates.** The predictive accuracy of S-map models was assessed across increasing forecast horizons (TP) and exclusion radii (ER), where ER was defined as TP+n (n={0,15}) to account for autocorrelation effects, for each target cyanobacterial group. The six best-performing parameter sets (selected via sequential leave-future-out cross-validation at a 1-day horizon) were used for evaluation. Model performance was quantified as Pearson's correlation coefficient rho (ρ) between observed and predicted values across forecast horizons from 1 to 30 days. Coloured points and lines represent the mean ρ of the best-performing S-map models and error bars show the standard deviation. The black line represents the performance of a benchmark persistence model, which assumes that future values remain unchanged (see **materials and methods** for details). Notably, the S-map models successfully capture the underlying system dynamics beyond autocorrelation effects, as their predictive performance remains stable even at a 30-day forecast horizon.

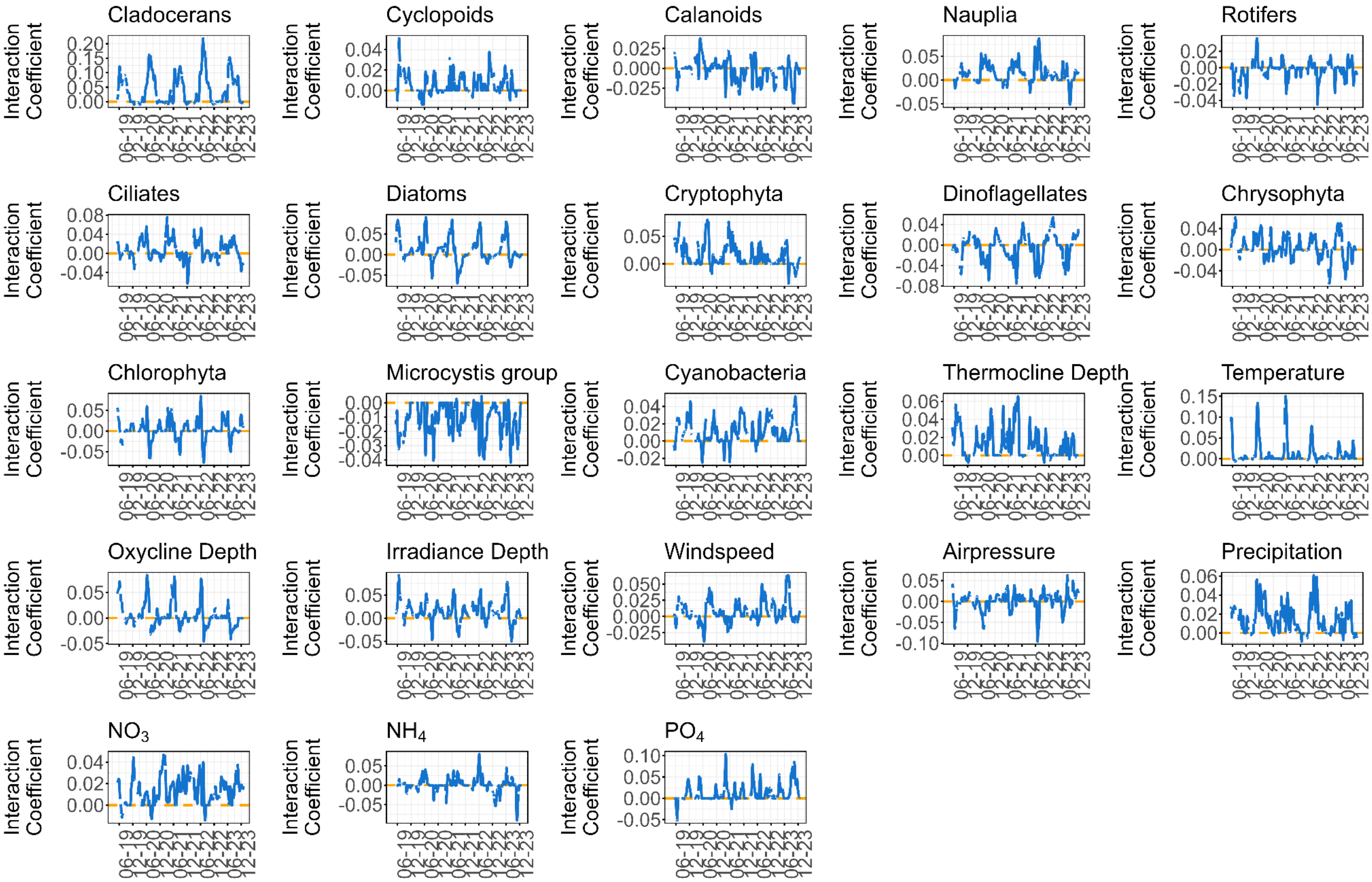


**Fig. S9: Time series of coefficients from the regularised multiview-distance S-maps (best model) for *Microcystis* group.**

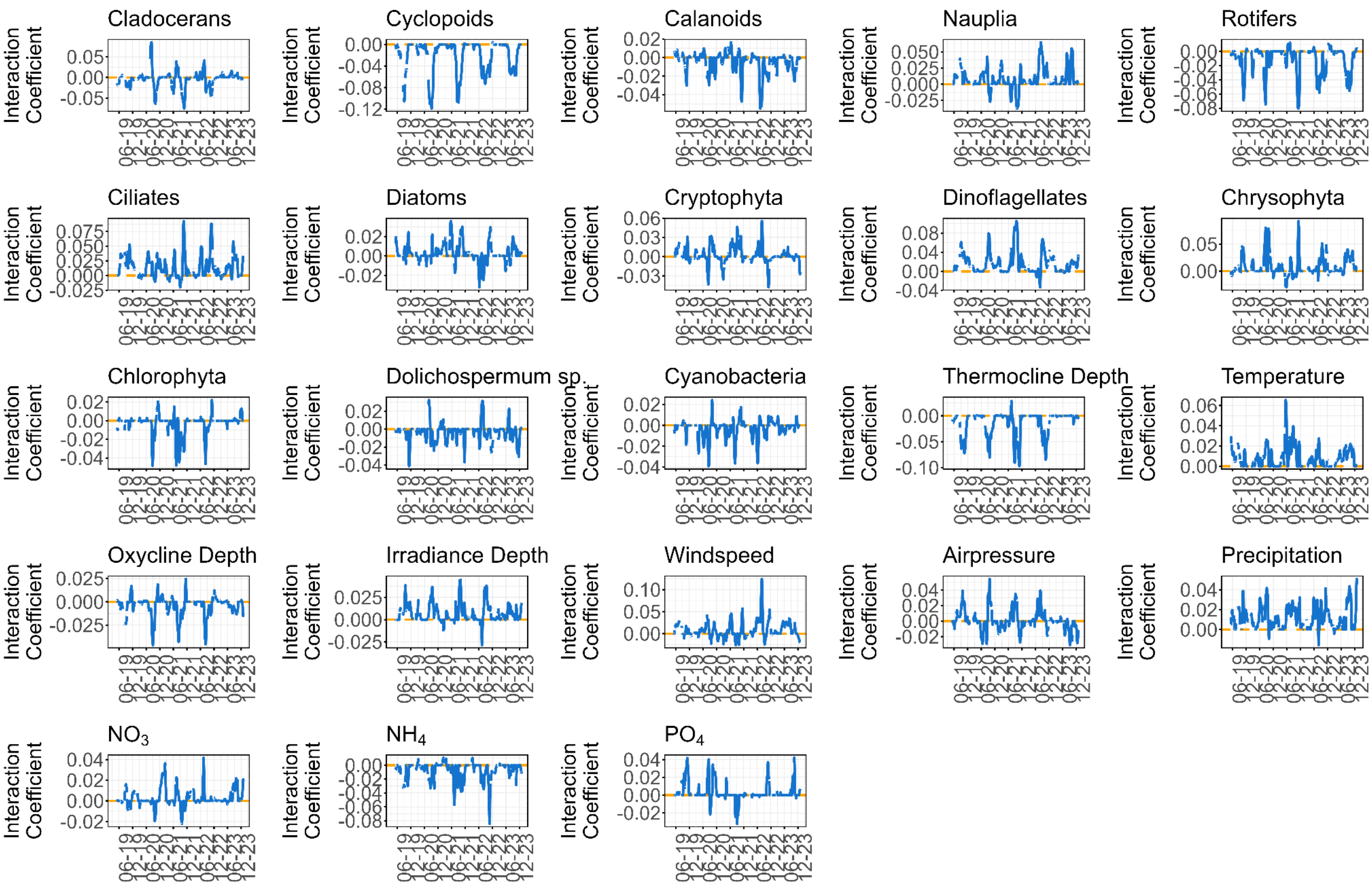


**Fig. S10: Time series of coefficients from the regularised multiview-distance S-maps (best model) for *Dolichospermum sp.***

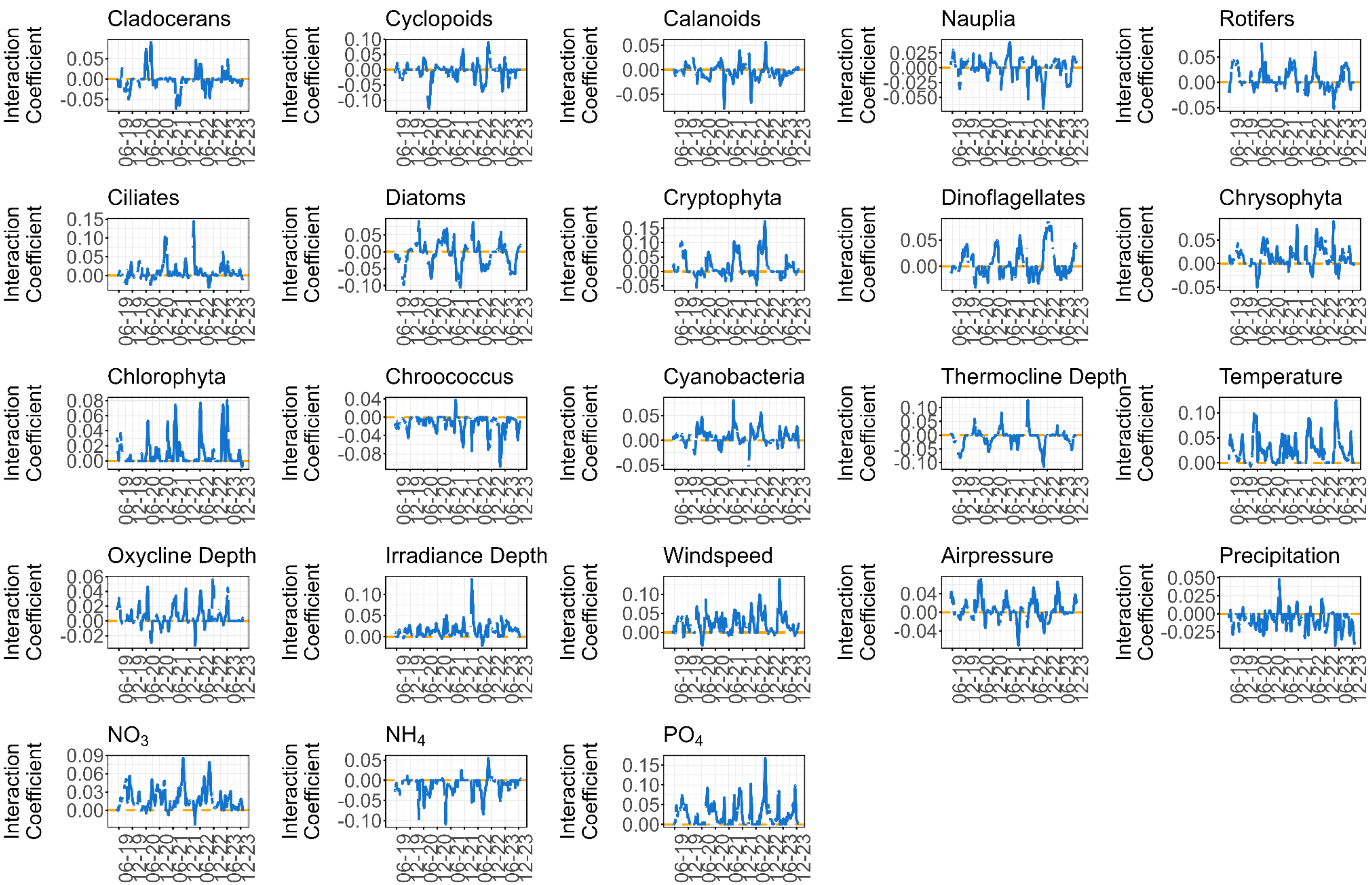


**Fig. S11: Time series of coefficients from the regularised multiview-distance S-maps (best model) for *Chroococcus sp.***

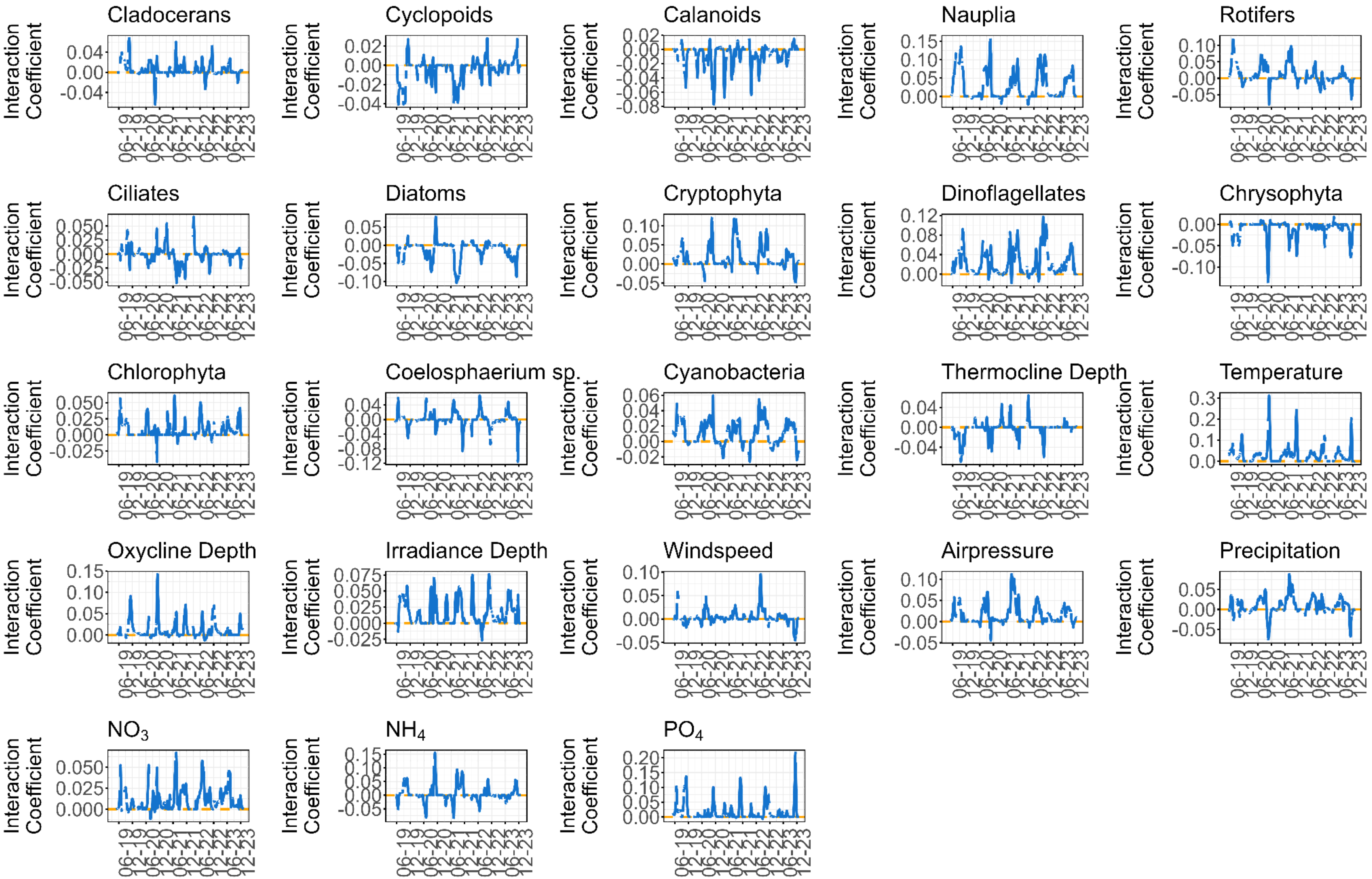


**Fig. S12: Time series of coefficients from the regularised multiview-distance S-maps (best model) for *Coelosphaerium sp.***

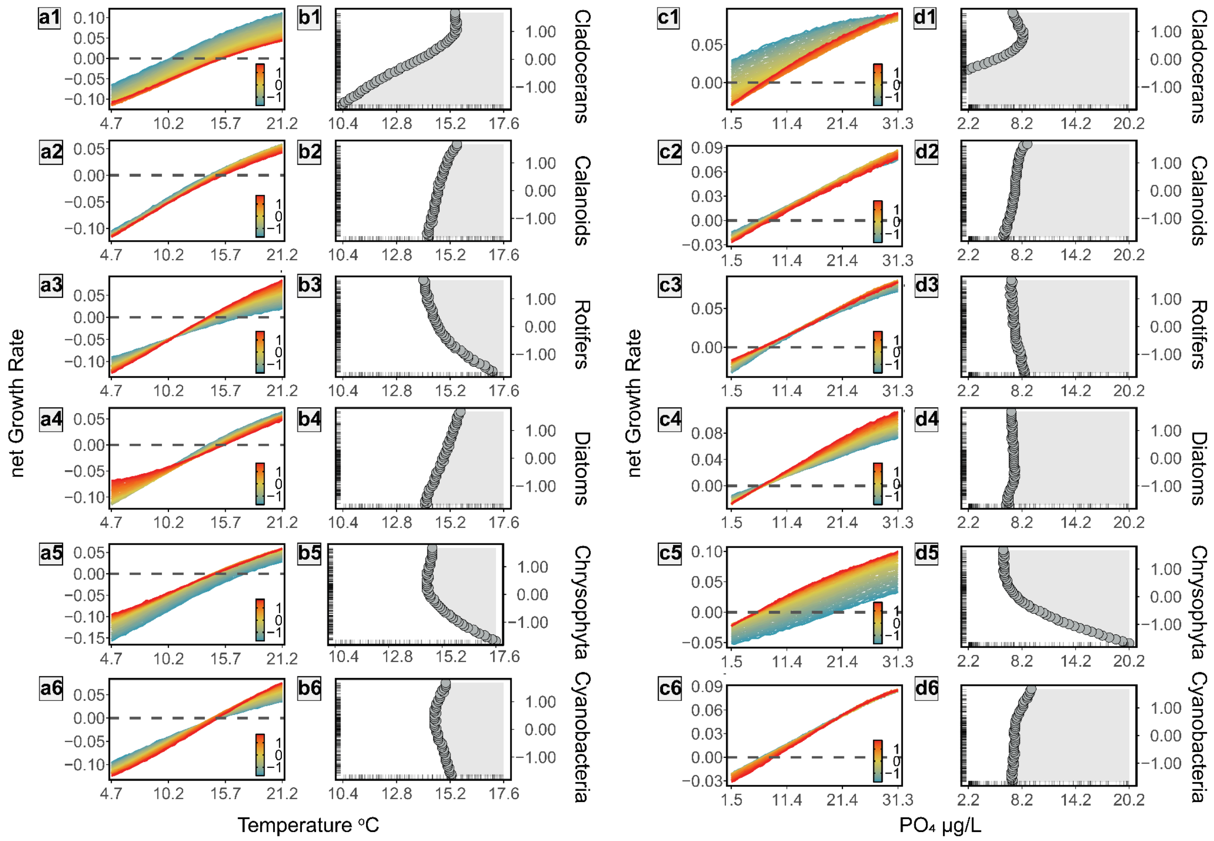


**Fig. S13: Biotic interactions shape *Chroococcus* sp. blooming across water temperature and phosphate gradients.** (**A1-A6**) Temperature dependency of *Chroococcus* nGR (Ln(ind/mL)·day$^{-1}$) across increasing densities (Ln(ind/mL)) of key planktonic groups: cladocerans (**1**), calanoids (**2**), rotifers (**3**), diatoms (**4**), chrysophytes (**5**), and other cyanobacteria (**6**). (**C1-C6**) Phosphate ($PO_4$) dependency of *Chroococcus* net growth rates across densities of the same planktonic groups. (**B1-B6, D1-D6**) Zero net growth analysis of blooming conditions along temperature and $PO_4$ gradients, respectively, as influenced by increasing densities of interacting organisms. Tick marks on the x- and y-axes indicate the empirical distributions of temperature, $PO_4$, and planktonic group scaled densities observed in Lake Greifensee from 2019 to 2023. Grey surfaces indicate conditions supporting positive net growth rates (blooming). See **Fig. 4** for a detailed interpretation of the relationships between blooming thresholds and organism densities. For detailed pairwise interaction analyses, see **Figs. S17 and S33**.

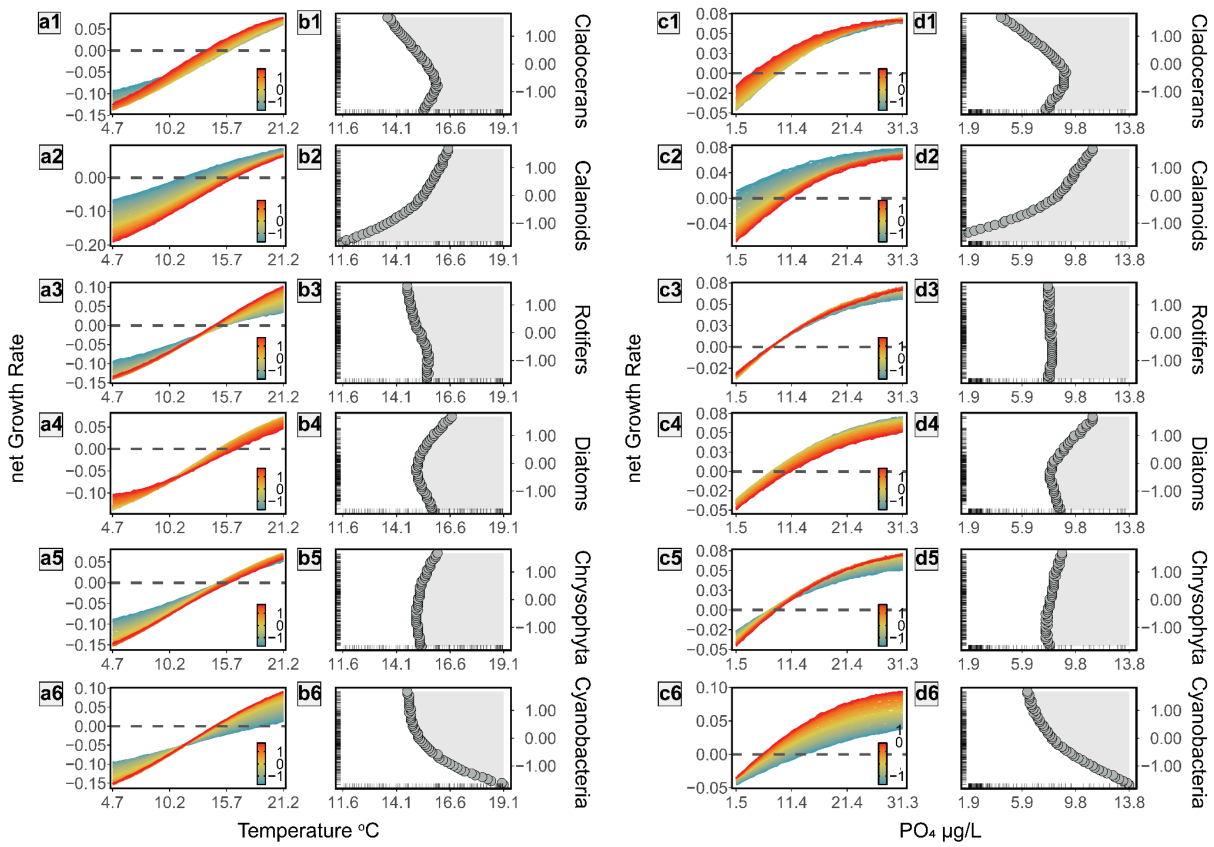


**Fig. S14: Biotic interactions shape *Coelosphaerium* sp. blooming across water temperature and phosphate gradients.** (**A1-A6**) Temperature dependency of *Coelosphaerium* nGR (Ln(ind/mL) $\cdot$ day$^{-1}$) across increasing densities (Ln(ind/mL)) of key planktonic groups: cladocerans (**1**), calanoids (**2**), rotifers (**3**), diatoms (**4**), chrysophytes (**5**), and other cyanobacteria (**6**). (**C1-C6**) Phosphate ($PO_4$) dependency of *Coelosphaerium* nGR across densities of the same planktonic groups. (**B1-B6, D1-D6**) Zero net growth analysis of blooming conditions along temperature and $PO_4$ gradients, respectively, as influenced by increasing densities of interacting organisms. Tick marks on the x- and y-axes indicate the empirical distributions of temperature, $PO_4$, and planktonic group scaled densities observed in Lake Greifensee from 2019 to 2023. Grey surfaces indicate conditions supporting positive net growth rates (blooming). See **Fig. 4** for a detailed interpretation of the relationships between blooming thresholds and organism densities. For detailed pairwise interaction analyses, see **Figs. S18 and S34**.

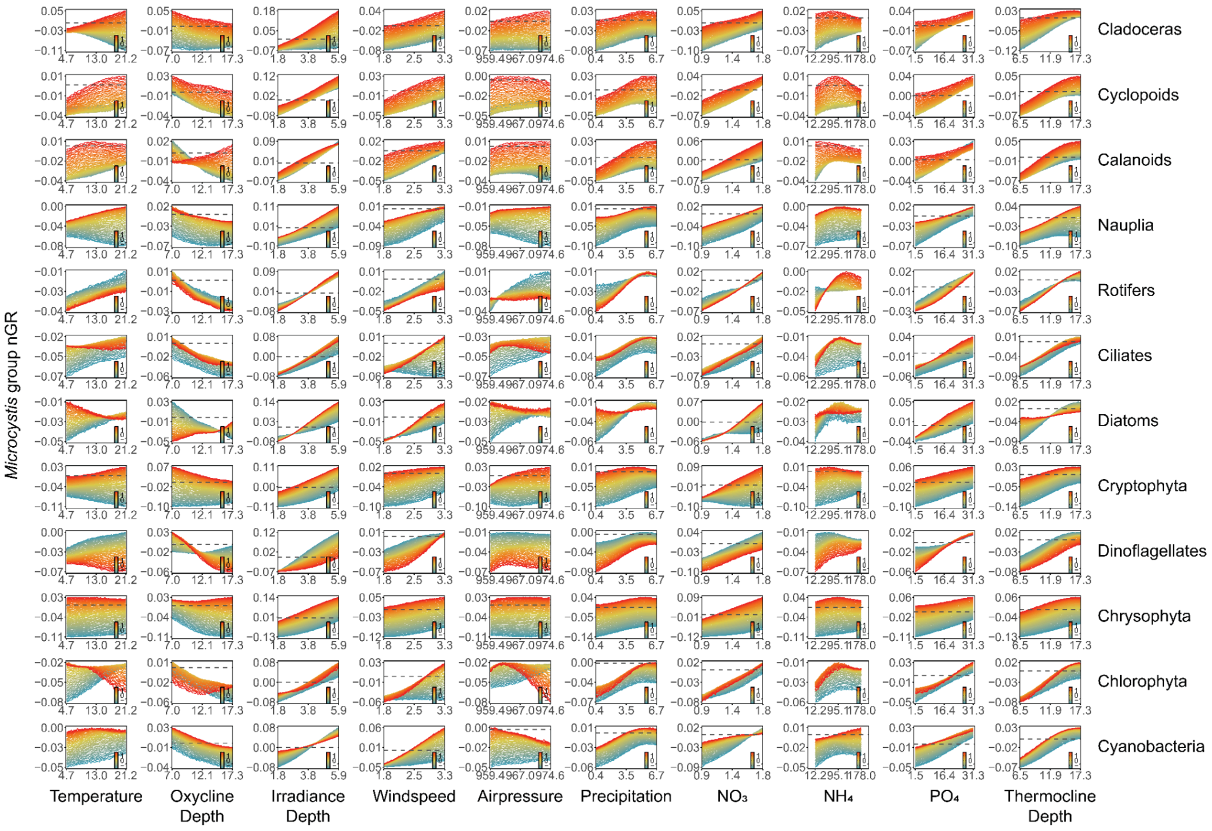


**Fig. S15: Pairwise interactions between biotic and abiotic factors shape *Microcystis* growth responses.** This figure shows how *Microcystis* nGRs (Ln(ind/mL) · day$^{-1}$) respond to combinations of environmental conditions, derived from regularized MDR S-maps analysis. Each panel shows predicted nGR when two variables (one biotic, one abiotic) vary across their observed range (5-95 percentile), while all other variables are held at median values. **Abiotic effects**: Positive correlations with most variables, except for a negative relationship with oxycline depth. **Biotic effects**: Mostly positive interactions, with notable negative effects from rotifers and dinoflagellates. **Interactive effects**: Biotic factors can modify or reverse growth responses to abiotic conditions, specifically: Temperature responses are modified by cladocerans, cyclopoid copepods, copepods' nauplii, ciliates, diatoms, cryptophytes, dinoflagellates and chlorophytes; Oxycline depth responses are altered by calanoid copepods, diatoms, cryptophytes, dinoflagellates, chrysophytes and chlorophytes; Air pressure responses are influenced by copepods' nauplii, rotifers, ciliates, diatoms, chlorophytes and cyanobacteria; $NH_4$ responses are modified by all biotic groups except chrysophytes and chlorophytes. **Units**: Biotic variables in Ln(ind/mL) and scaled; abiotic variables: temperature (°C), precipitation (mm/day), wind speed (m/s), $PO_4$ and $NH_4$ (µg/L), $NO_3$ (mg/L), air pressure (hPa), depth measurements (metres).

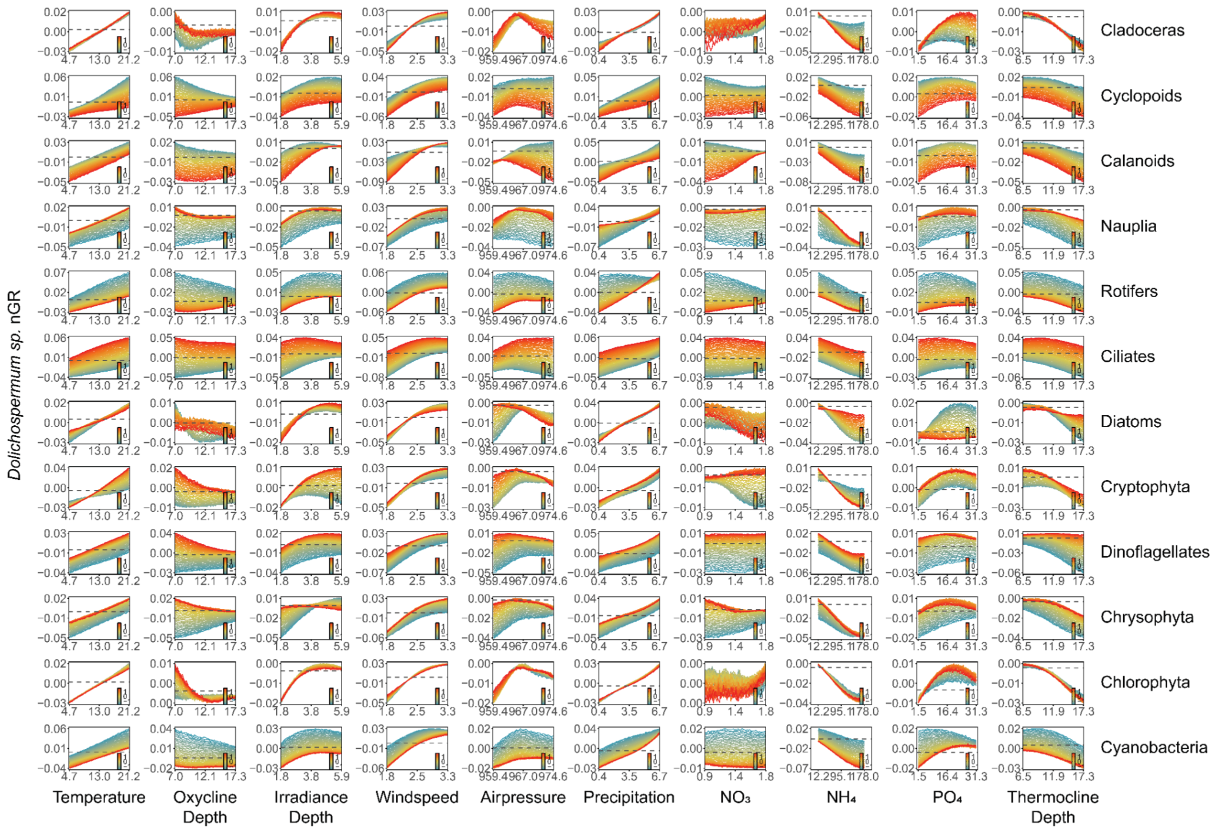


**Fig. S16: Pairwise interactions between biotic and abiotic factors shape *Dolichospermum* sp. growth responses.** This figure shows how *Dolichospermum* sp. nGRs (Ln(ind/mL)·day$^{-1}$) respond to combinations of environmental conditions, derived from regularized MDR S-maps analysis (as in **Fig. S15**). Each panel shows predicted nGR when two variables (one biotic, one abiotic) vary across their observed range (5-95 percentile), while all other variables are held at median values. **Abiotic effects**: Positive correlations with temperature, light irradiance, wind speed, precipitation, and $PO_4$; negative relationships with oxycline depth, $NH_4$, and thermocline depth; unimodal response to air pressure; negligible effects of $NO_3$. **Biotic effects**: Negative effects from cyclopoid and calanoid copepods, rotifers, and cyanobacteria; positive effects from nauplii, ciliates, dinoflagellates, and chrysophytes. **Interactive effects**: Biotic factors can modify or reverse growth responses to abiotic conditions, specifically: Oxycline depth responses are modified by cyclopoid copepods, rotifers, diatoms, dinoflagellates, chrysophytes, and chlorophytes; $NO_3$ responses are altered by cladocerans, calanoid copepods, rotifers, diatoms, cryptophytes, and chrysophytes; $PO_4$ responses are influenced by rotifers and diatoms. **Units** as in **Fig. S15**.

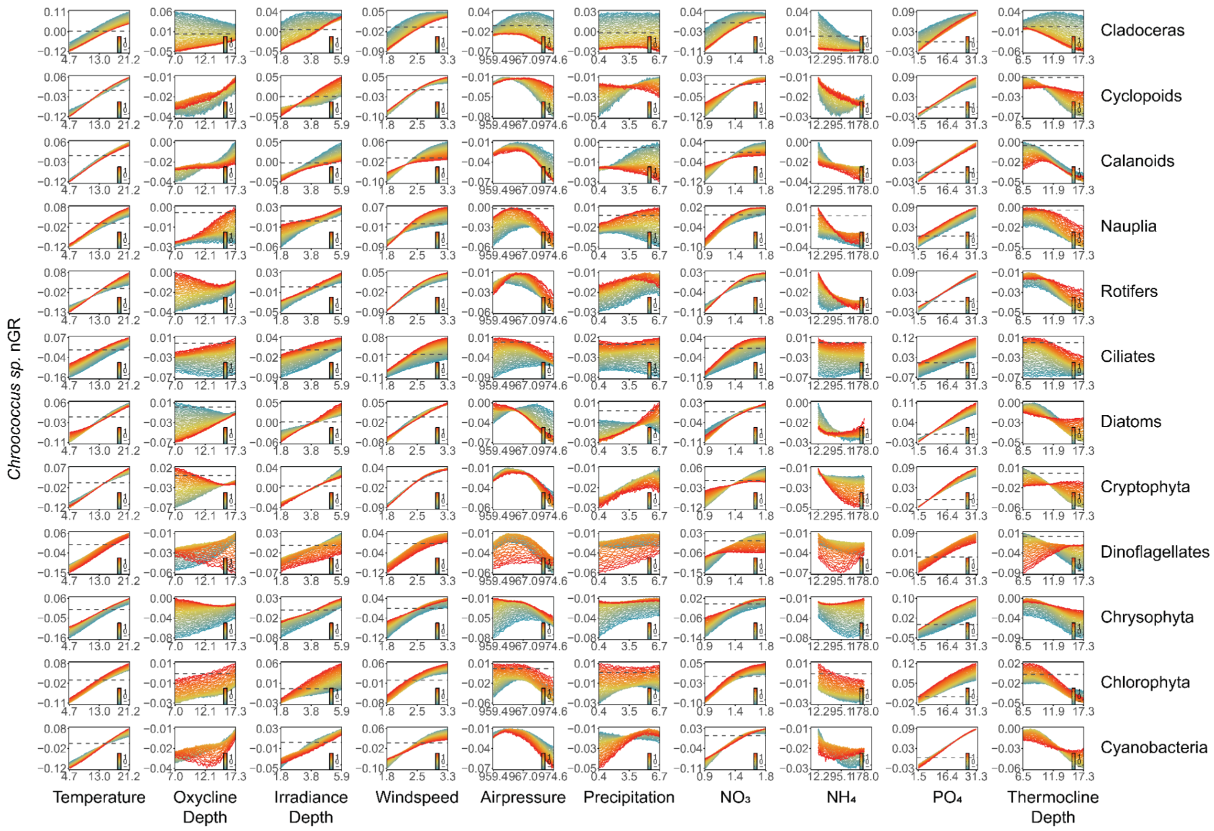


**Fig. S17: Pairwise interactions between biotic and abiotic factors shape *Chroococcus* sp. growth responses.** This figure shows how *Chroococcus* sp. nGRs (Ln(ind/mL)·day$^{-1}$) respond to combinations of environmental conditions, derived from regularized MDR S-maps analysis (as in **Fig. S15**). Each panel shows predicted nGR when two variables (one biotic, one abiotic) vary across their observed range (5-95 percentile), while all other variables are held at median values. **Abiotic effects**: Positive correlations with temperature, oxycline depth, light irradiance, wind speed, $NO_3$, and $PO_4$; negative relationships with $NH_4$ and thermocline depth; unimodal response to air pressure; negligible effects of precipitation. **Biotic effects**: Negative effects from cladocerans, cyclopoid copepods, and diatoms; positive effects from calanoid copepods, their nauplii, rotifers, ciliates, chrysophytes, and chlorophytes. **Interactive effects**: Biotic factors can modify or reverse growth responses to abiotic conditions, specifically: Oxycline depth responses are modified by most biotic groups except cyclopoid copepods and chlorophytes; Precipitation responses are altered by copepods (cyclopoid, calanoid, nauplii), diatoms, dinoflagellates, chrysophytes, chlorophytes, and cyanobacteria; $PO_4$ responses are influenced by cladocerans, ciliates, diatoms, cryptophytes, and dinoflagellates. **Units** as in **Fig. S15**.

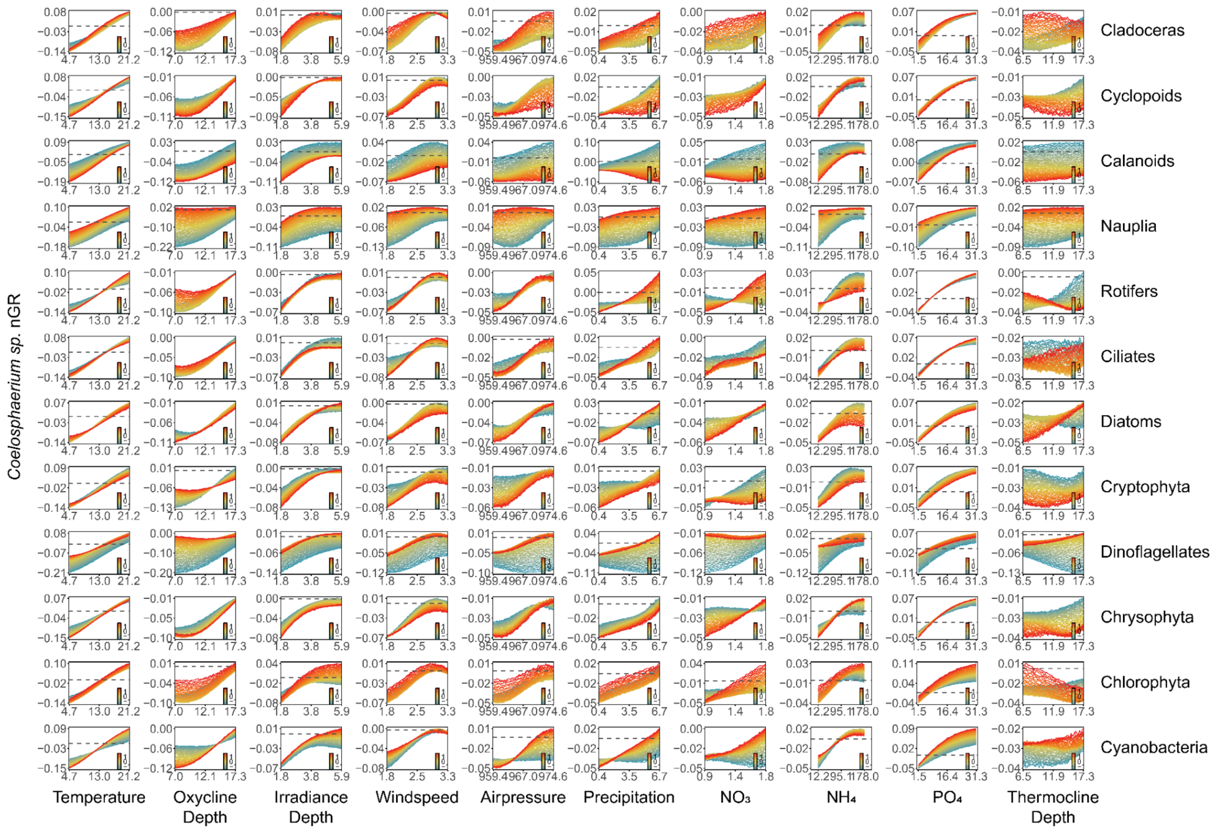


**Fig. S18: Pairwise interactions between biotic and abiotic factors shape *Coelosphaerium* sp. growth responses.** This figure shows how *Coelosphaerium* sp. nGRs (Ln(ind/mL) · day$^{-1}$) respond to combinations of environmental conditions, derived from regularized MDR S-maps analysis (as in **Fig. S15**). Each panel shows predicted nGR when two variables (one biotic, one abiotic) vary across their observed range (5-95 percentile), while all other variables are held at median values. **Abiotic effects**: Positive correlations with all abiotic variables. **Biotic effects**: Negative effects from cyclopoid and calanoid copepods, cryptophytes, and chrysophytes; positive effects from cladocerans, copepods, dinoflagellates, and cyanobacteria. **Interactive effects**: Biotic factors can modify or reverse growth responses to abiotic conditions, specifically: Air pressure, precipitation, and $NO_3$ responses are modified by calanoid copepods and their nauplii, diatoms, cryptophytes, dinoflagellates, and cyanobacteria; Thermocline depth responses are altered by cladocerans, cyclopoid copepods, rotifers, diatoms, dinoflagellates, and chlorophytes. **Units** as in **Fig. S15**.

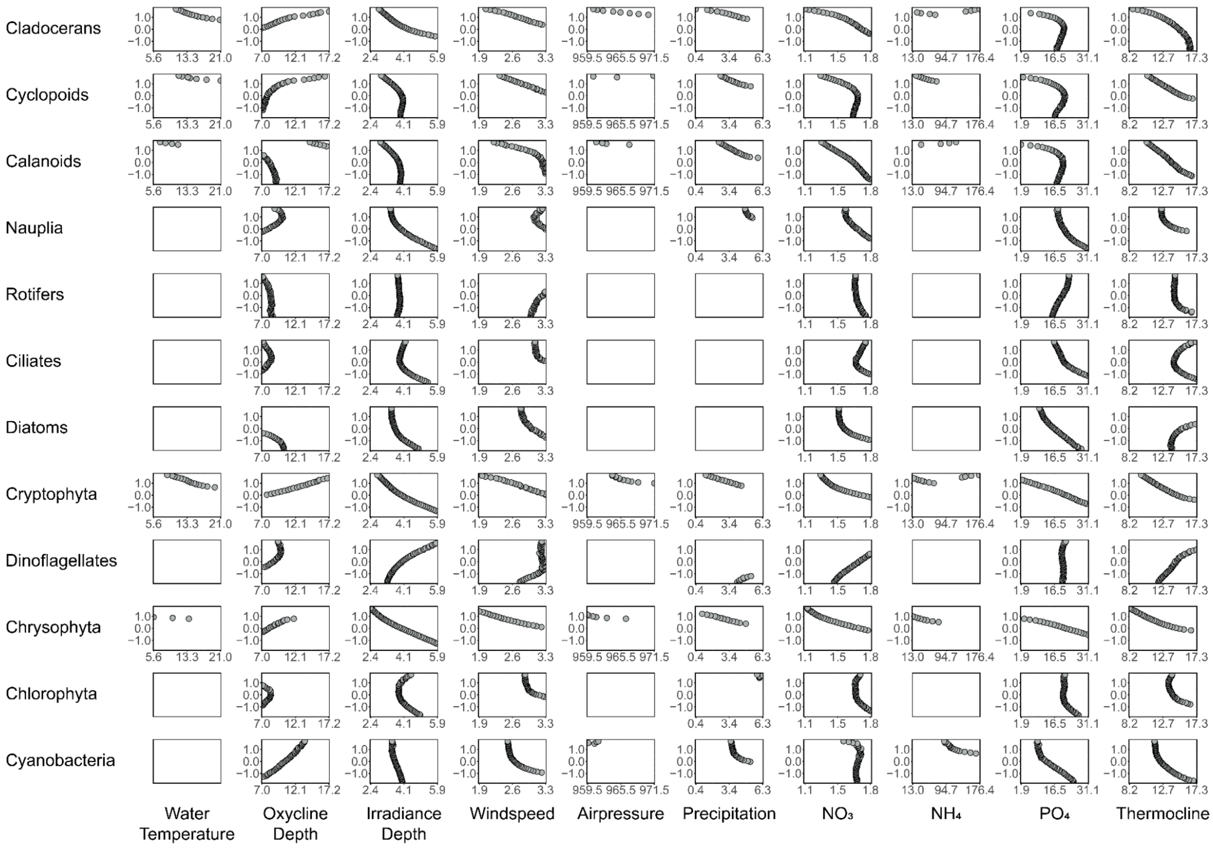


**Fig. S19: Zero net growth analysis for *Microcystis* bloom formation across biotic-abiotic variable combinations.** This figure analyzes bloom initiation conditions by identifying zero-net growth isoclines from the response functions in **Fig. S15**. Each panel shows how the minimum environmental conditions required for blooming change with increasing densities of different planktonic groups. **Response patterns**: Positive correlations between blooming points and organism density indicate bloom inhibition (higher environmental thresholds needed); negative correlations indicate bloom facilitation (lower thresholds needed). **Key influences**: Cladocerans, cryptophytes, and chrysophytes strongly affect blooming conditions across all tested abiotic ranges. **Notable absences**: White panels indicate conditions where blooming was never observed due to consistently negative or positive growth rates. **Units** as in **Fig. S15**.

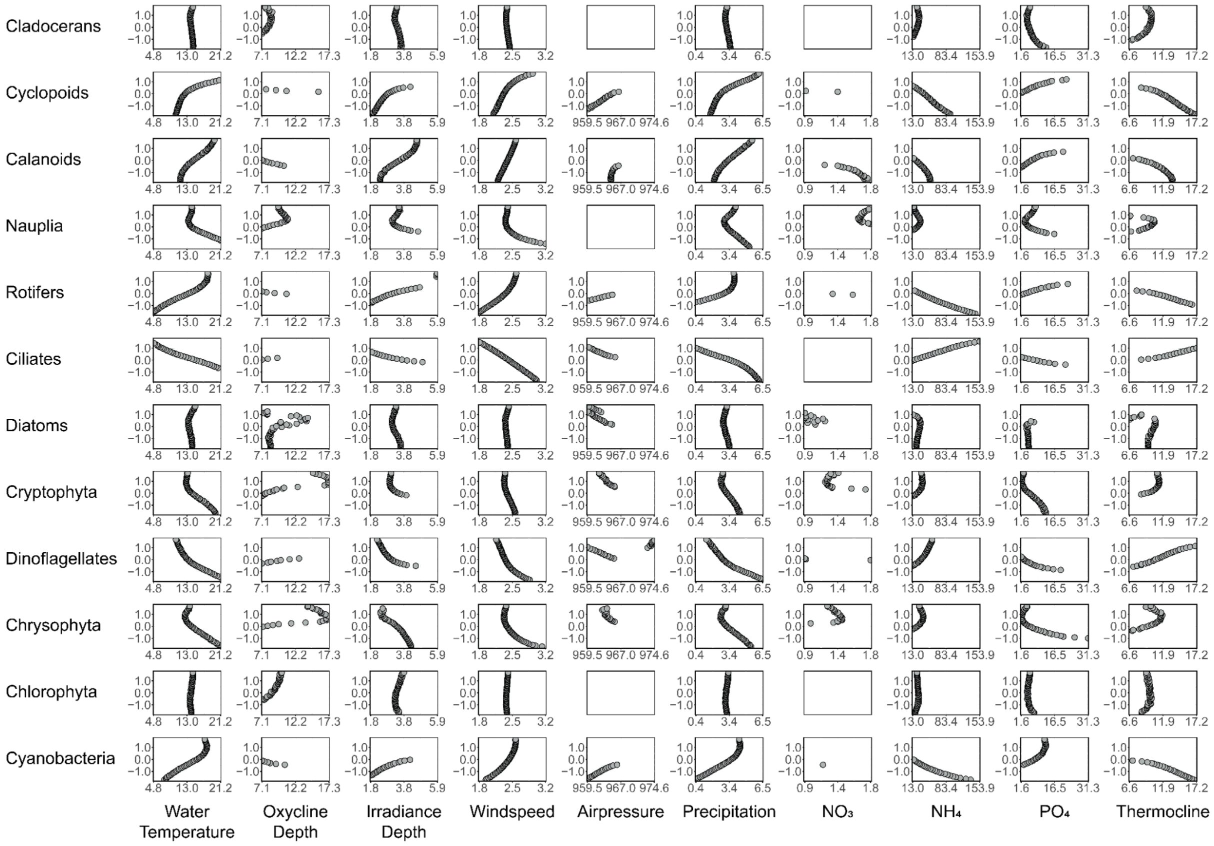


**Fig. S20: Zero net growth analysis for *Dolichospermum* bloom formation across biotic-abiotic variable combinations.** This figure analyzes bloom initiation conditions by identifying zero-net growth isoclines from the response functions in **Fig. S16**. Each panel shows how the minimum environmental conditions required for blooming change with increasing densities of different planktonic groups. **Response patterns**: Positive correlations between blooming points and organism density indicate bloom inhibition (higher environmental thresholds needed); negative correlations indicate bloom facilitation (lower thresholds needed). **Key influences**: Rotifers, ciliates, dinoflagellates, and cyanobacteria strongly affect blooming conditions across most abiotic scenarios and throughout their observed density ranges. **Notable absences**: White panels indicate conditions where blooming was never observed due to consistently negative or positive growth rates. **Units** as in **Fig. S15**.

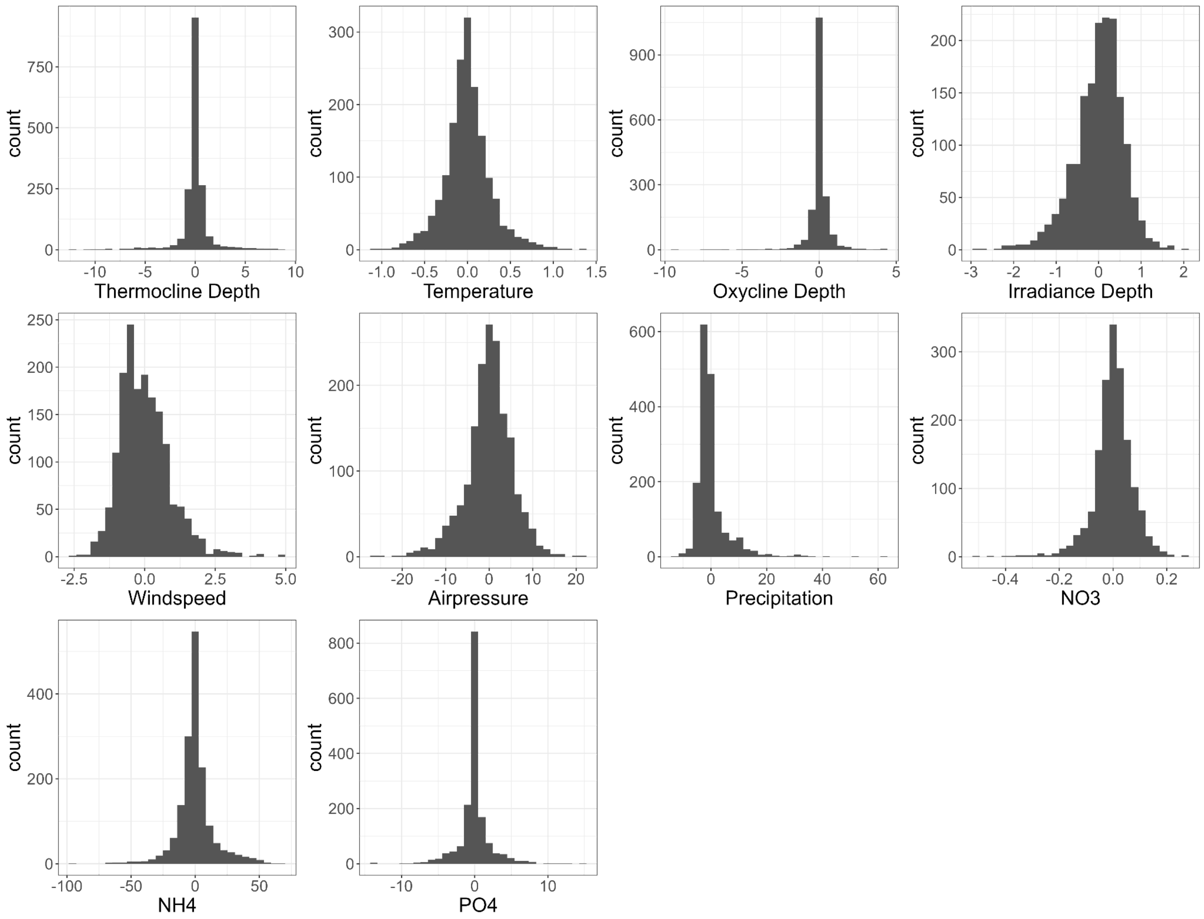


**Fig. S21: Histograms of the residuals obtained after smoothing the abiotic variables using a loess function (span = 0.03) to reduce random noise in the raw data.**

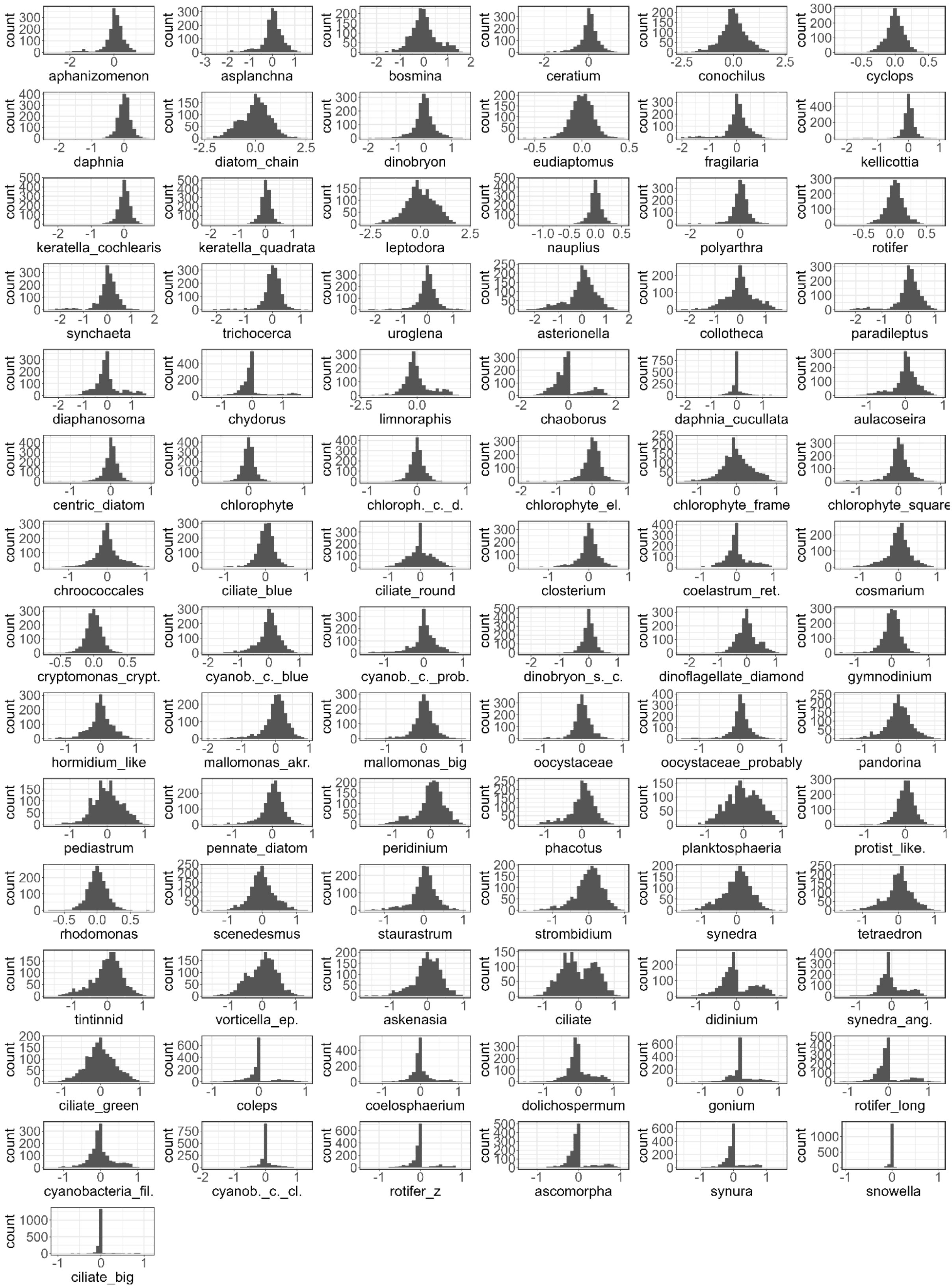


**Fig. S22: Histograms of the residuals obtained after smoothing the biotic variables using a loess function (span = 0.03) to reduce random noise in the raw data.** The biotic variables represent individual taxa before their aggregation into broader taxonomic groups (see **Tables S1 and S2** for the groupings).

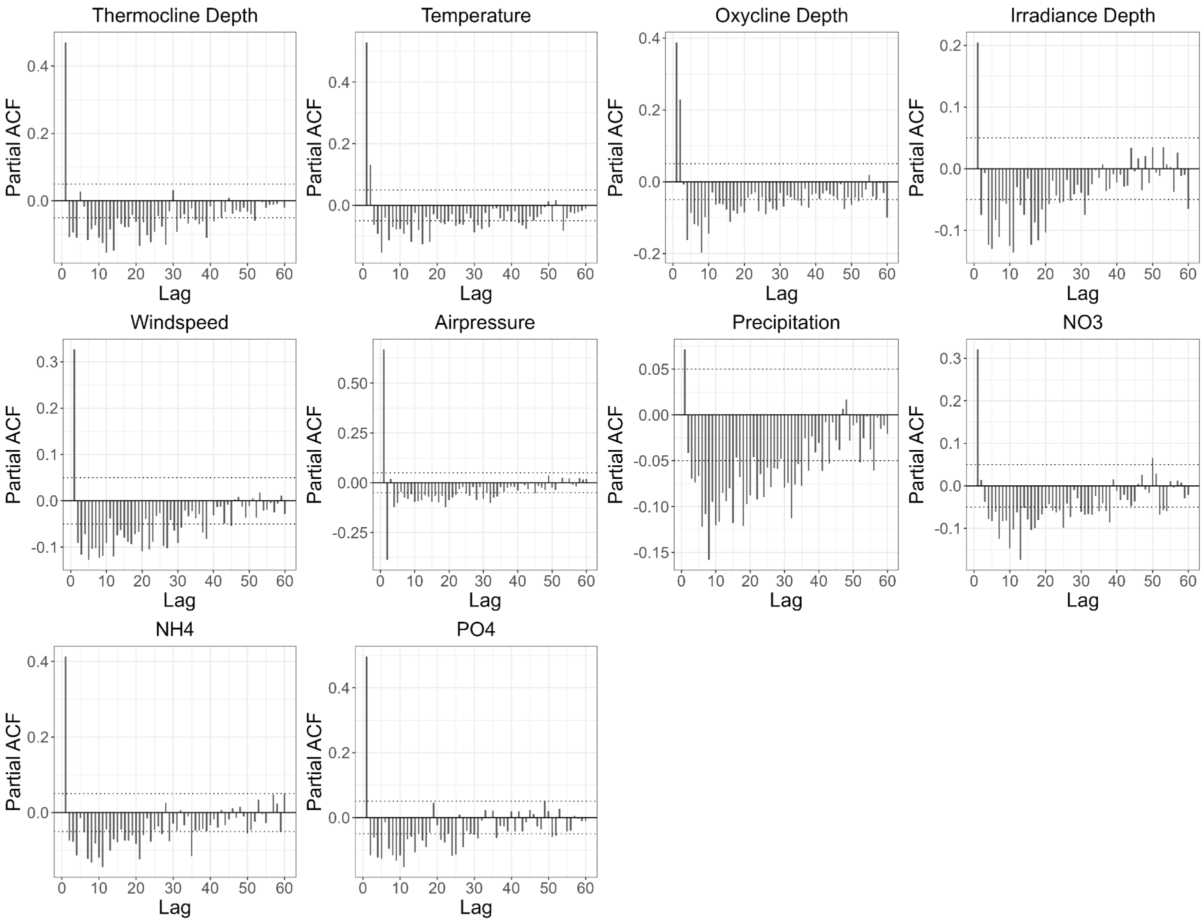


**Fig. S23: Partial autocorrelation plots of the residuals obtained after smoothing the abiotic variables using a loess function (span = 0.03) to reduce random noise in the raw data.**

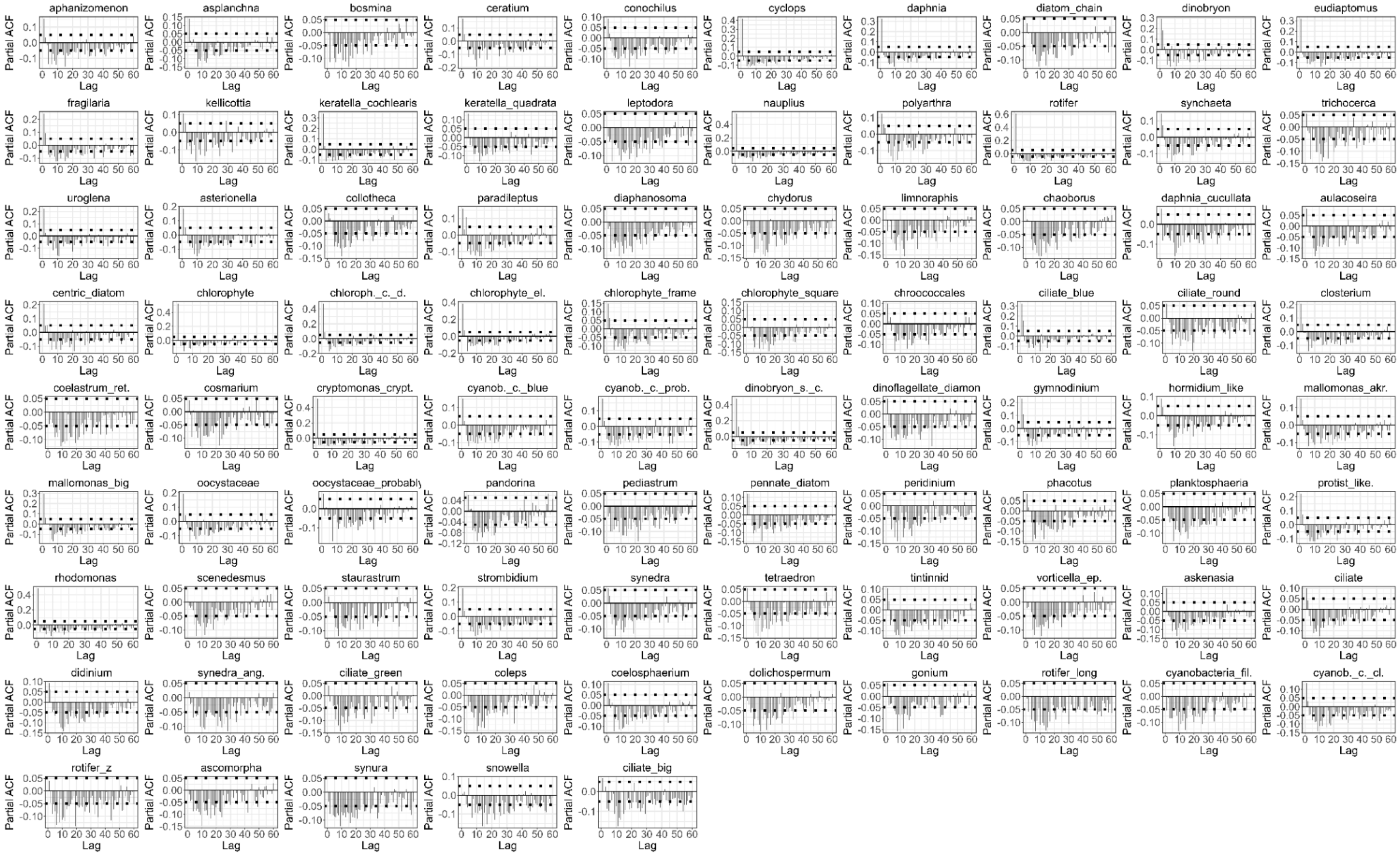

**Fig. S24: Partial autocorrelation plots of the residuals obtained after smoothing the biotic variables using a loess function (span = 0.03) to reduce random noise in the raw data.** The biotic variables represent individual taxa before their aggregation into broader taxonomic groups (see **Tables S1 and S2** for the groupings).

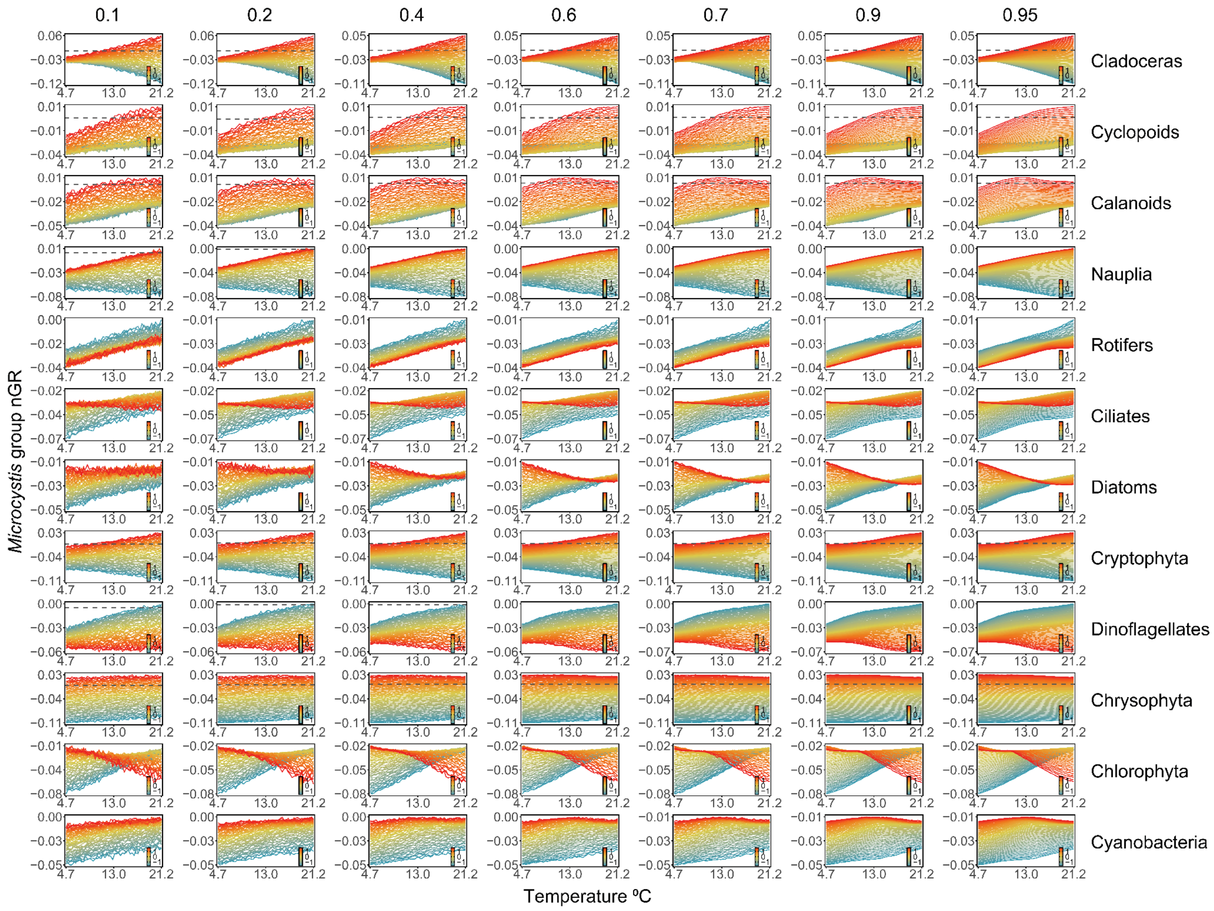


**Fig. S25: Sensitivity analysis of *Microcystis* growth to temperature across different dataset sizes.** Response functions of *Microcystis* group net growth rate (nGR) (Ln(ind/mL) · day$^{-1}$) to temperature across different biotic interactions (rows) and data subset sizes (columns). Each panel represents a biotic interaction, with temperature on the x-axis and changes in nGR on the y-axis. Columns correspond to different proportions of the dataset used (10%–95%) to assess response function robustness. Lines show mean estimated responses from 50 subsamples (see **materials and methods**) and colour indicates the scaled density of the biotic factor in Ln(ind/mL). Patterns remain consistent across data lengths, confirming robustness.

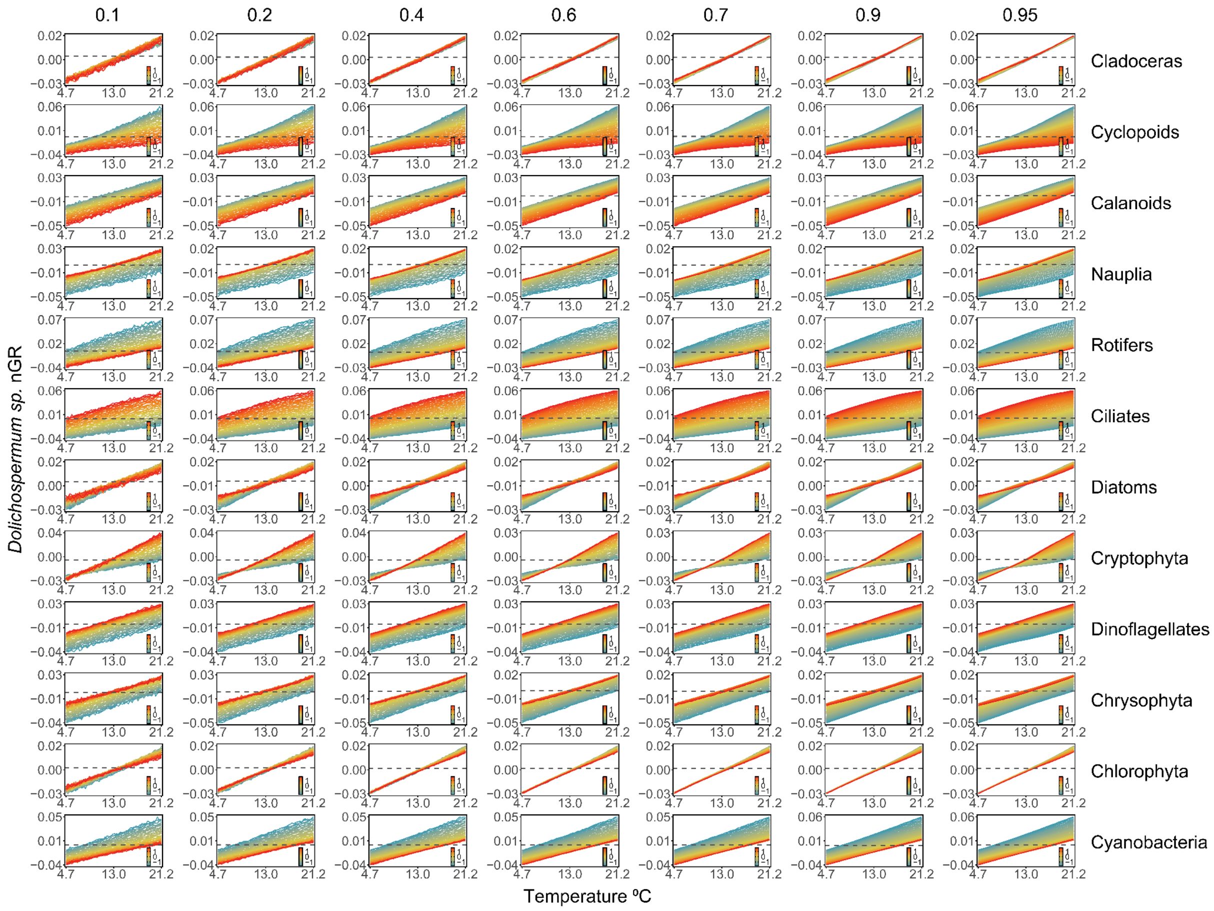


**Fig. S26: Sensitivity analysis of *Dolichospermum* growth to temperature across different dataset sizes.** Response functions of *Dolichospermum sp.* net growth rate (nGR) (Ln(ind/mL)·day$^{-1}$) to temperature across different biotic interactions (rows) and data subset sizes (columns). Each panel represents a biotic interaction, with temperature on the x-axis and changes in nGR on the y-axis. Columns correspond to different proportions of the dataset used (10%–95%) to assess response function robustness. Lines show mean estimated responses from 50 subsamples (see **materials and methods**) and colour indicates the scaled density of the biotic factor in Ln(ind/mL). Patterns remain consistent across data lengths, confirming robustness.

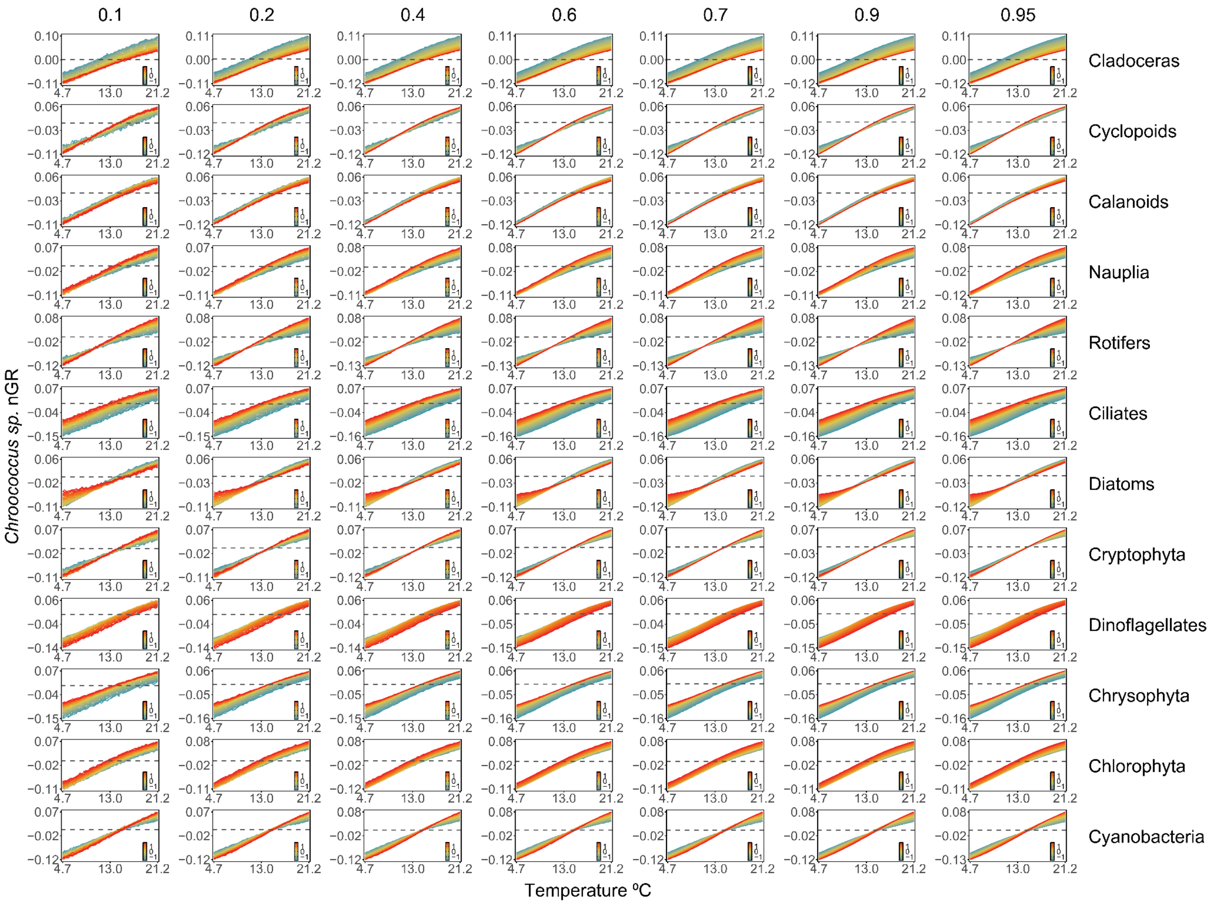


**Fig. S27: Sensitivity analysis of *Chroococcus* growth to temperature across different dataset sizes.** Response functions of *Chroococcus sp.* net growth rate (nGR) (Ln(ind/mL) · $day^{-1}$) to temperature across different biotic interactions (rows) and data subset sizes (columns). Each panel represents a biotic interaction, with temperature on the x-axis and changes in nGR on the y-axis. Columns correspond to different proportions of the dataset used (10%–95%) to assess response function robustness. Lines show mean estimated responses from 50 subsamples (see **materials and methods**) and colour indicates the scaled density of the biotic factor in Ln(ind/mL). Patterns remain consistent across data lengths, confirming robustness.

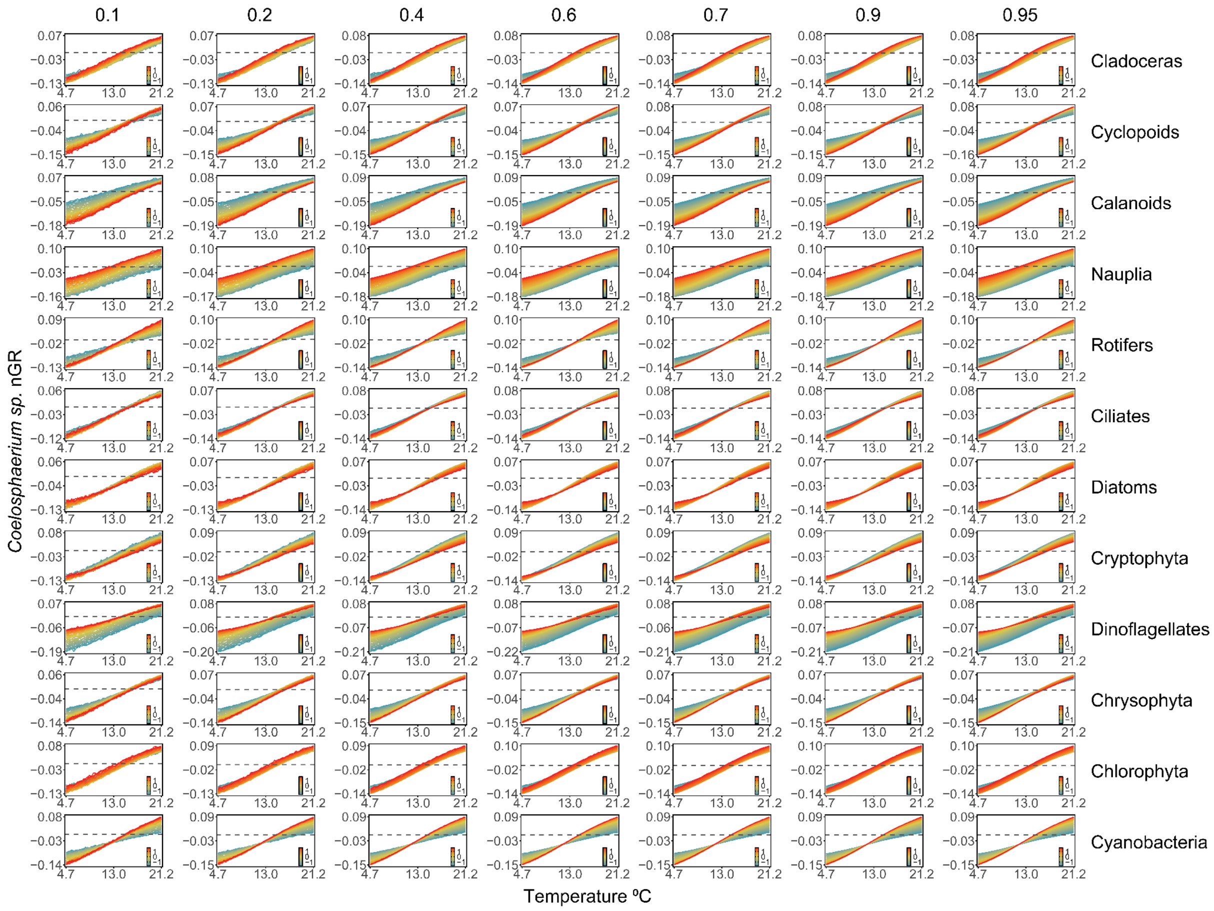


**Fig. S28: Sensitivity analysis of *Coelosphaerium* growth to temperature across different dataset sizes.** Response functions of *Coelosphaerium sp.* net growth rate (nGR) (Ln(ind/mL) · $day^{-1}$) to temperature across different biotic interactions (rows) and data subset sizes (columns). Each panel represents a biotic interaction, with temperature on the x-axis and changes in nGR on the y-axis. Columns correspond to different proportions of the dataset used (10%–95%) to assess response function robustness. Lines show mean estimated responses from 50 subsamples (see **materials and methods**) and colour indicates the scaled density of the biotic factor in Ln(ind/mL). Patterns remain consistent across data lengths, confirming robustness.

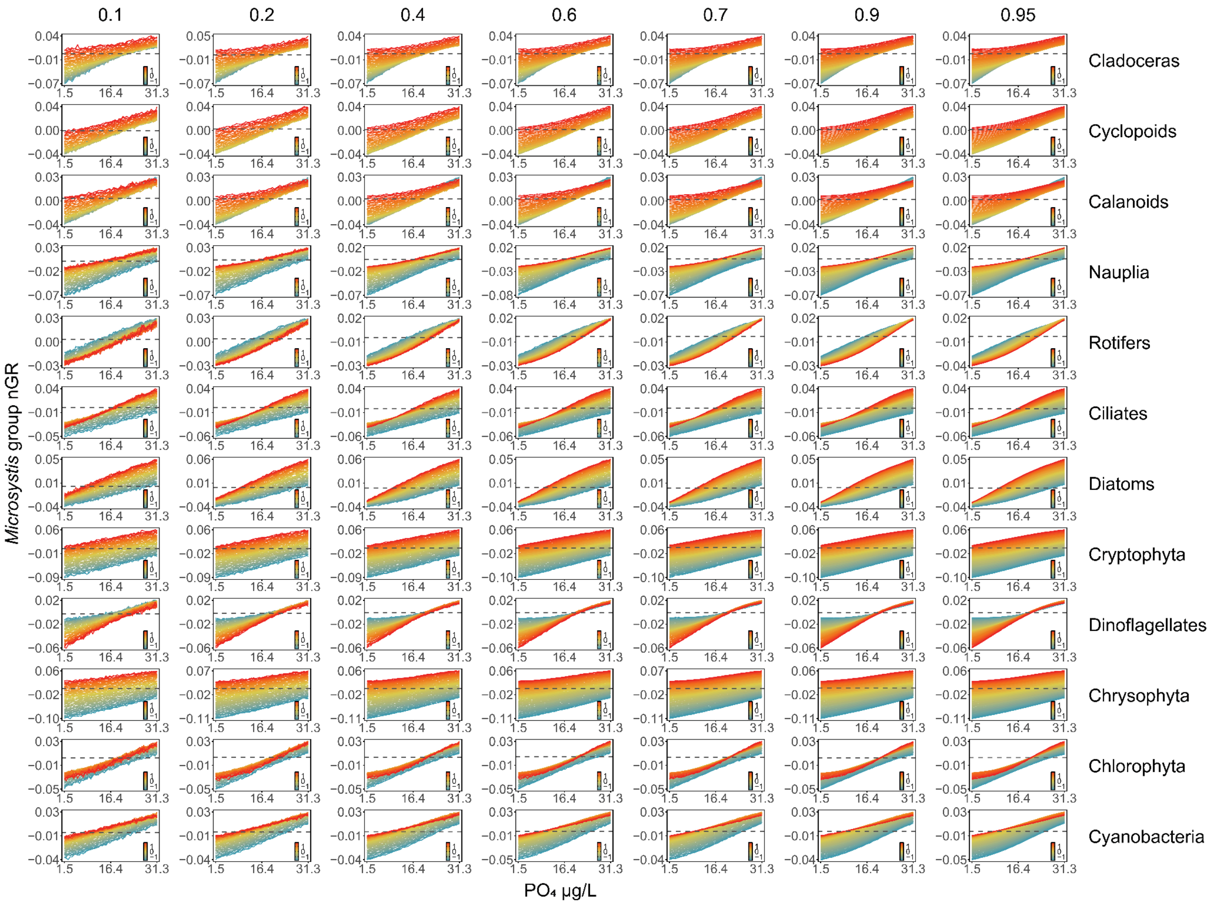


**Fig. S29: Sensitivity analysis of *Microcystis* growth to phosphate across different dataset sizes.** Response functions of *Microcystis* group net growth rate (nGR) (Ln(ind/mL) · day$^{-1}$) to $PO_4$ across different biotic interactions (rows) and data subset sizes (columns). Each panel represents a biotic interaction, with $PO_4$ on the x-axis and changes in nGR on the y-axis. Columns correspond to different proportions of the dataset used (10%–95%) to assess response function robustness. Lines show mean estimated responses from 50 subsamples (see **materials and methods**) and colour indicates the scaled density of the biotic factor in Ln(ind/mL). Patterns remain consistent across data lengths, confirming robustness.

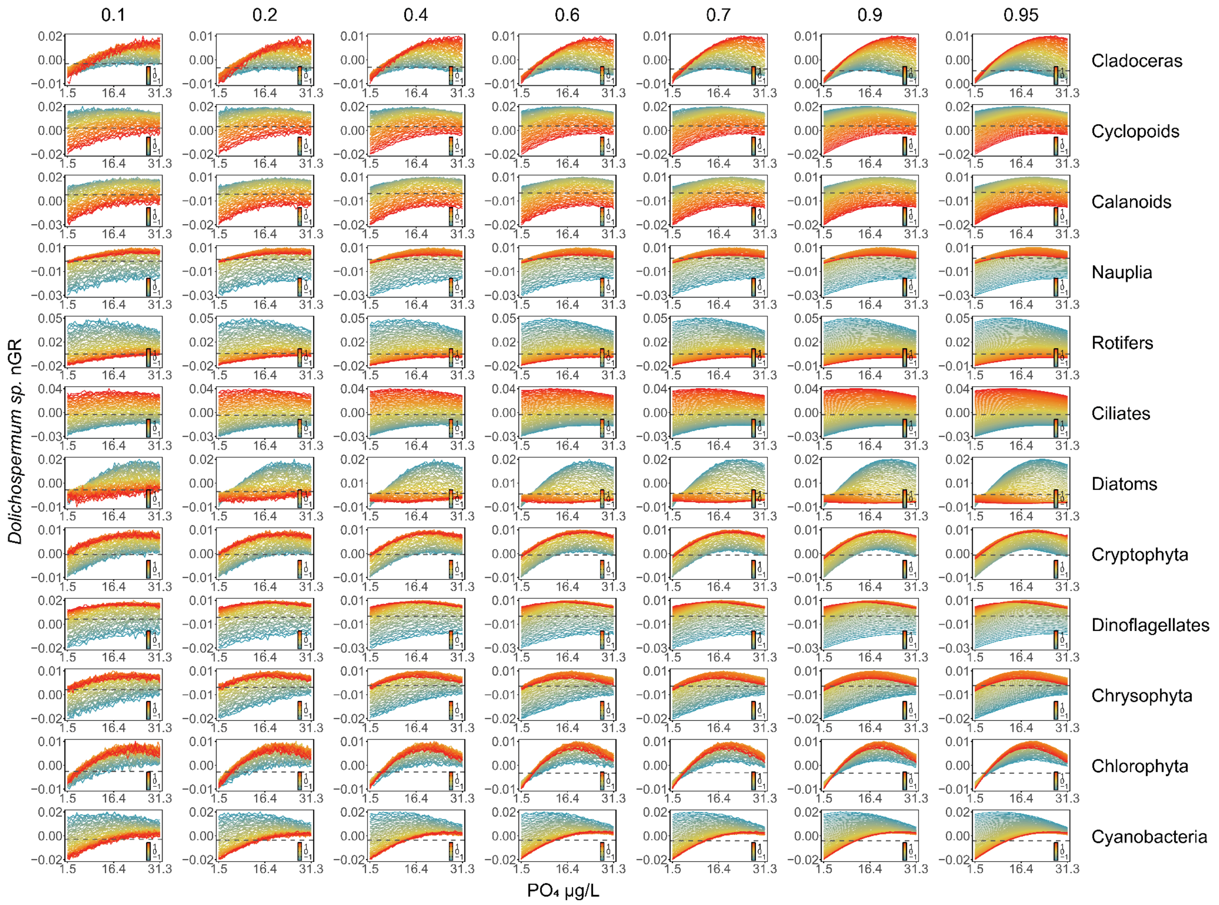


**Fig. S30: Sensitivity analysis of *Dolichospermum* growth to phosphate across different dataset sizes.** Response functions of *Dolichospermum sp.* net growth rate (nGR) (Ln(ind/mL)·day$^{-1}$) to $PO_4$ across different biotic interactions (rows) and data subset sizes (columns). Each panel represents a biotic interaction, with $PO_4$ on the x-axis and changes in nGR on the y-axis. Columns correspond to different proportions of the dataset used (10%–95%) to assess response function robustness. Lines show mean estimated responses from 50 subsamples (see **materials and methods**) and colour indicates the scaled density of the biotic factor in Ln(ind/mL). Patterns remain consistent across data lengths, confirming robustness.

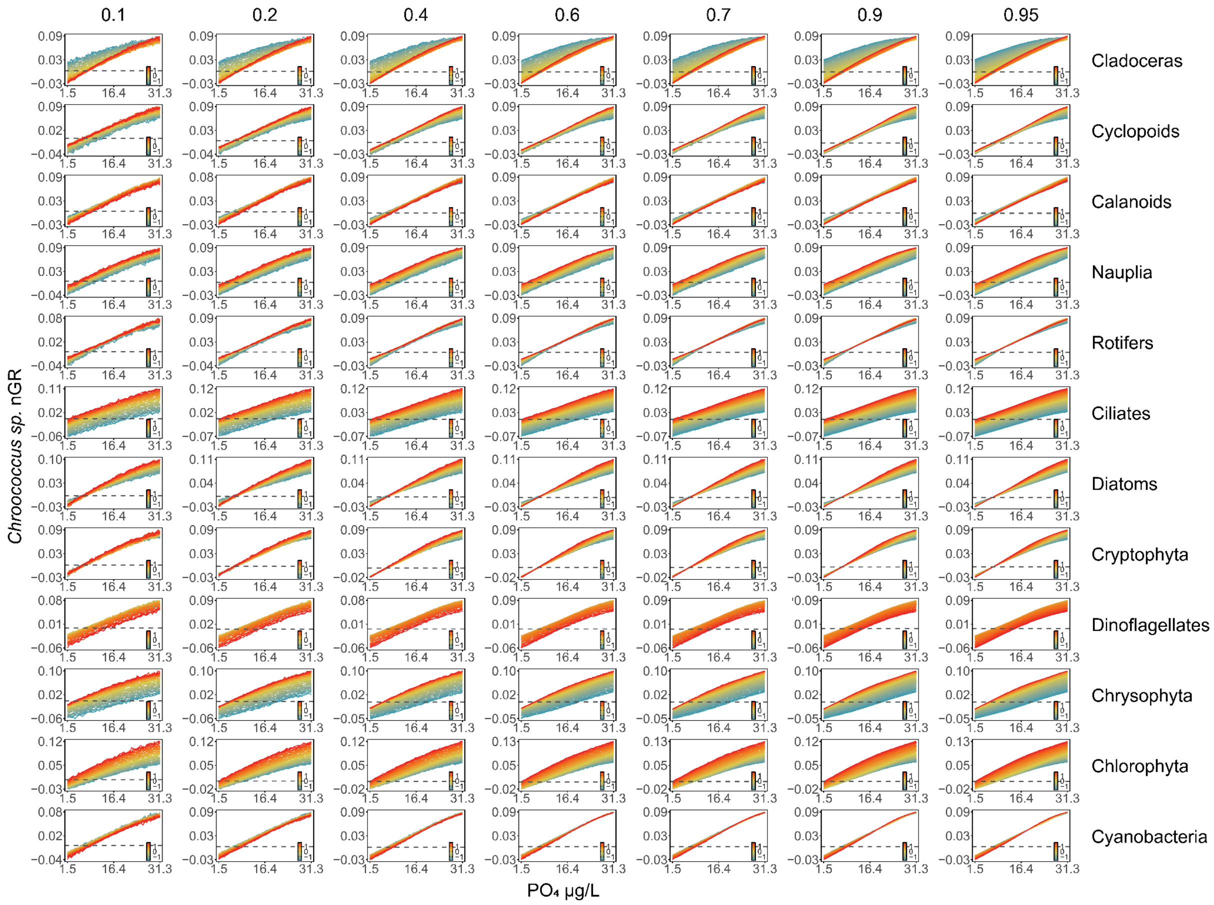


**Fig. S31: Sensitivity analysis of *Chroococcus* growth to phosphate across different dataset sizes.** Response functions of *Chroococcus sp.* net growth rate (nGR) (Ln(ind/mL) · day$^{-1}$) to $PO_4$ across different biotic interactions (rows) and data subset sizes (columns). Each panel represents a biotic interaction, with $PO_4$ on the x-axis and changes in nGR on the y-axis. Columns correspond to different proportions of the dataset used (10%–95%) to assess response function robustness. Lines show mean estimated responses from 50 subsamples (see **materials and methods**) and colour indicates the scaled density of the biotic factor in Ln(ind/mL). Patterns remain consistent across data lengths, confirming robustness.

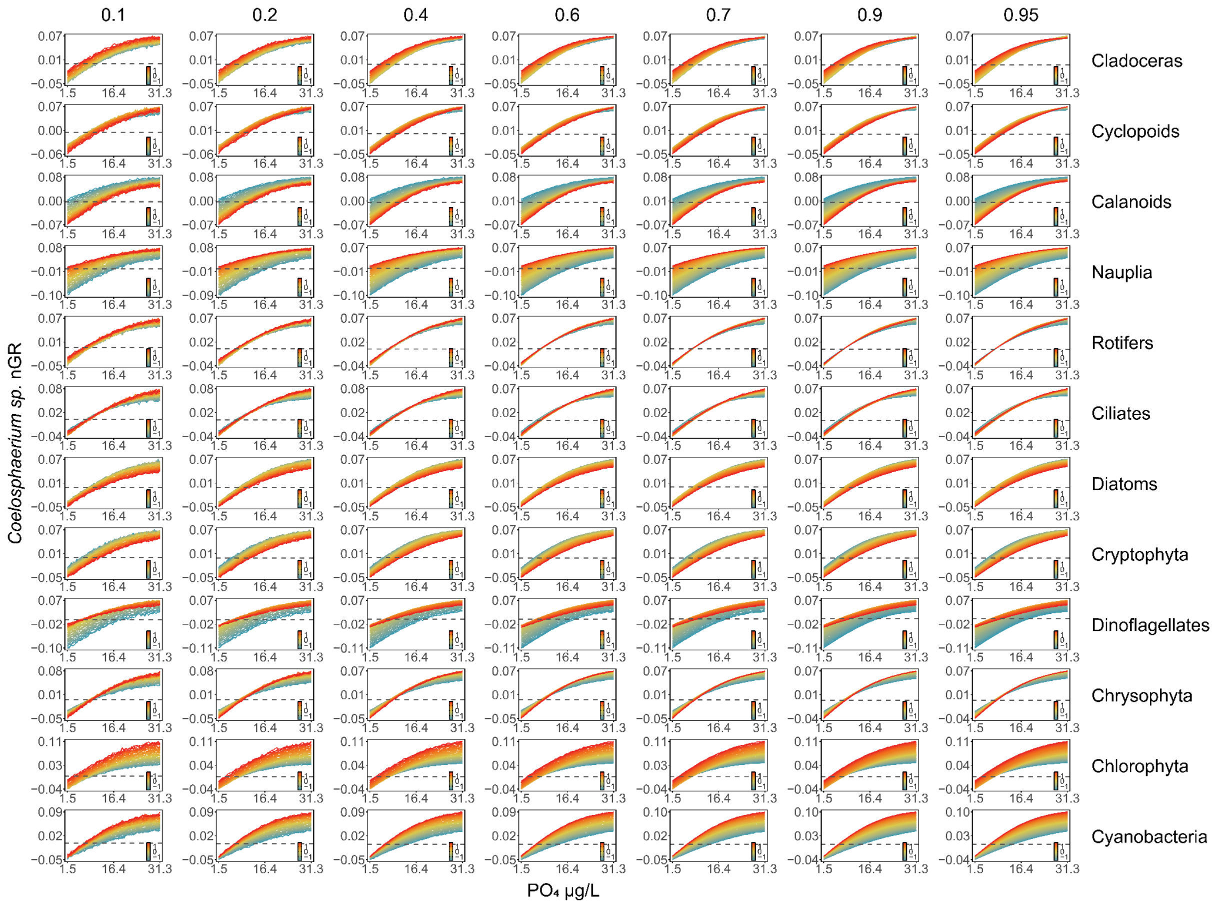


**Fig. S32: Sensitivity analysis of *Coelosphaerium* growth to phosphate across different dataset sizes.** Response functions of *Coelosphaerium sp*. net growth rate (nGR) (Ln(ind/mL)·day$^{-1}$) to $PO_4$ across different biotic interactions (rows) and data subset sizes (columns). Each panel represents a biotic interaction, with $PO_4$ on the x-axis and changes in nGR on the y-axis. Columns correspond to different proportions of the dataset used (10%–95%) to assess response function robustness. Lines show mean estimated responses from 50 subsamples (see **materials and methods**) and colour indicates the scaled density of the biotic factor in Ln(ind/mL). Patterns remain consistent across data lengths, confirming robustness.

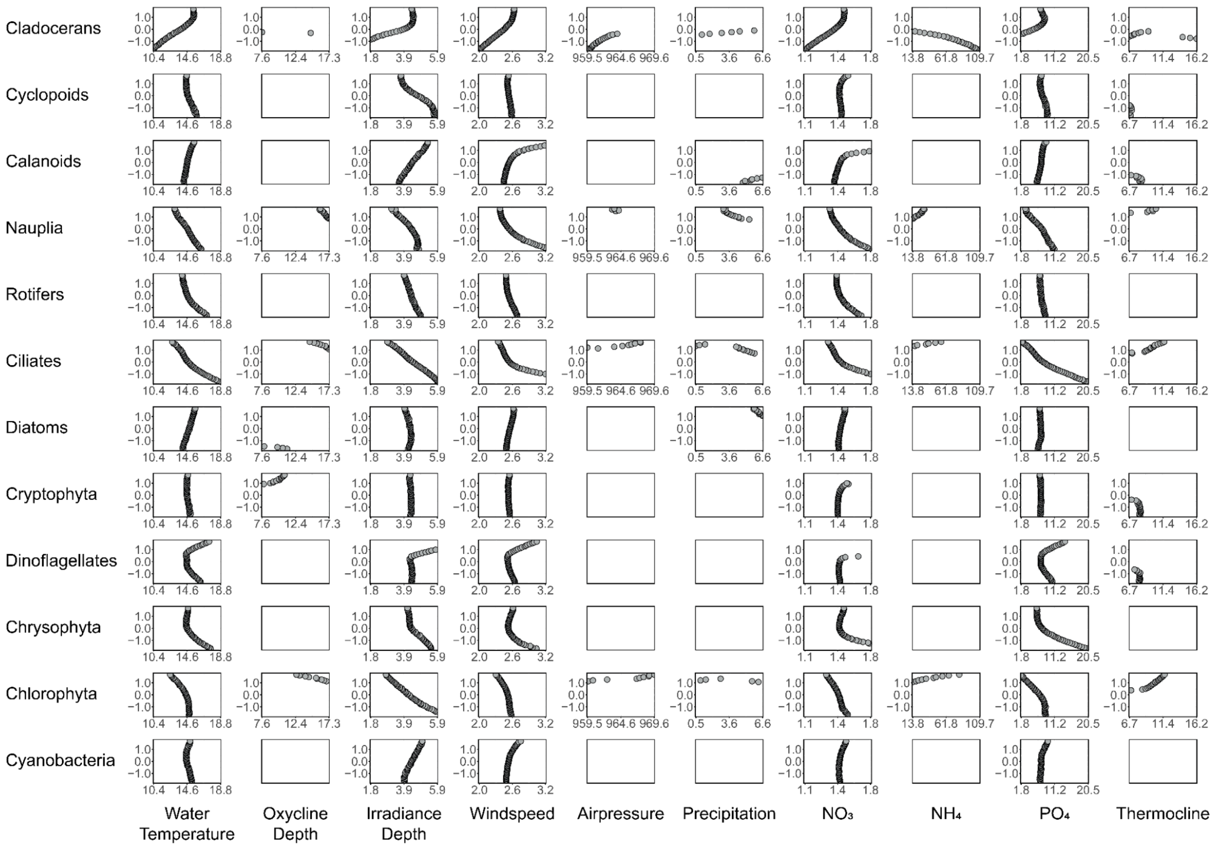


**Fig. S33: Zero net growth analysis for *Chroococcus* bloom formation across biotic-abiotic variable combinations.** This figure analyzes bloom initiation conditions by identifying zero-net growth isoclines from the response functions in **Fig. S17**. Each panel shows how the minimum environmental conditions required for blooming change with increasing densities of different planktonic groups. **Response patterns**: Positive correlations between blooming points and organism density indicate bloom inhibition (higher environmental thresholds needed); negative correlations indicate bloom facilitation (lower thresholds needed). **Key influences**: Cladocerans and ciliates strongly affect blooming conditions across most abiotic scenarios. **Notable absences**: White panels indicate conditions where blooming was never observed due to consistently negative or positive growth rates. **Units** as in **Fig. S15**.

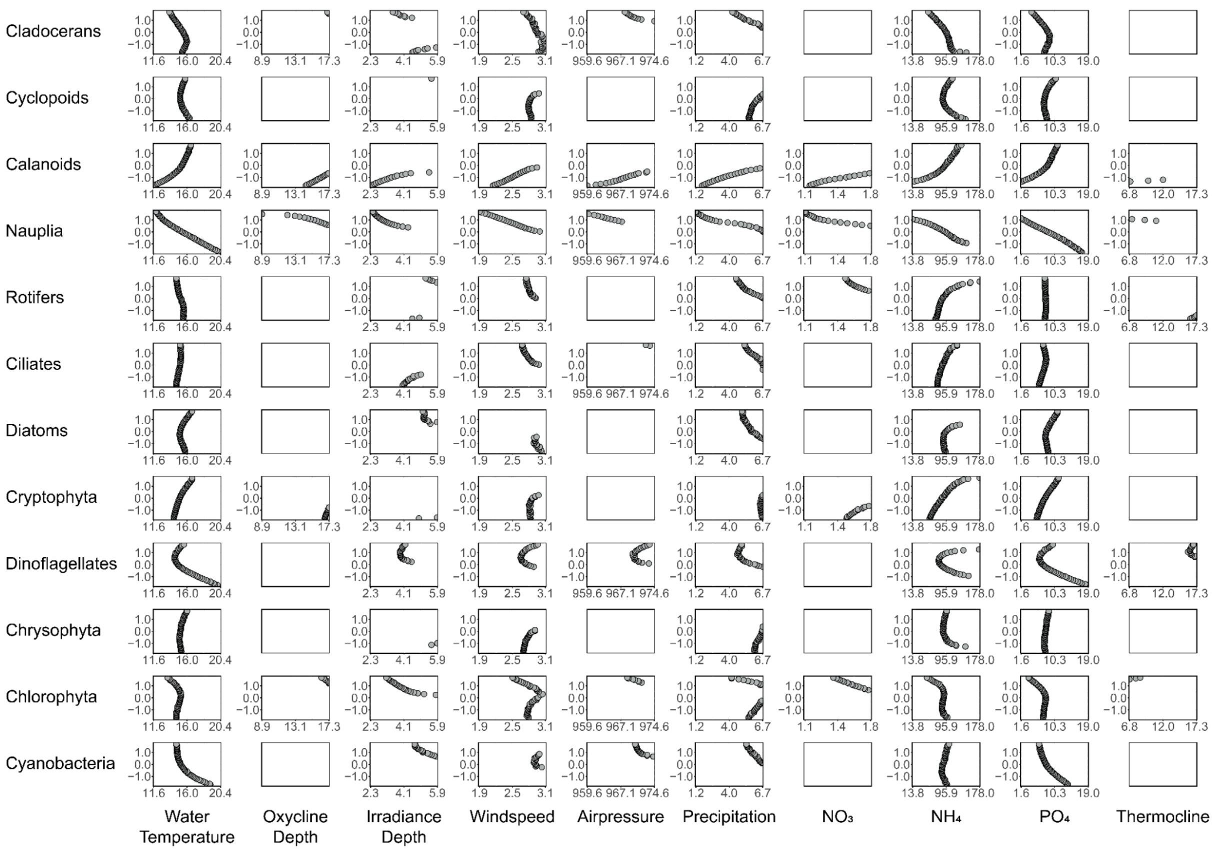


**Fig. S34: Zero net growth analysis for *Coelosphaerium* bloom formation across biotic-abiotic variable combinations.** This figure analyzes bloom initiation conditions by identifying zero-net growth isoclines from the response functions in **Fig. S18**. Each panel shows how the minimum environmental conditions required for blooming change with increasing densities of different planktonic groups. **Response patterns**: Positive correlations between blooming points and organism density indicate bloom inhibition (higher environmental thresholds needed); negative correlations indicate bloom facilitation (lower thresholds needed). **Key influences**: Calanoid copepods and their nauplii strongly affect blooming conditions across most abiotic scenarios. **Notable absences**: White panels indicate conditions where blooming was never observed due to consistently negative or positive growth rates. **Units** as in **Fig. S15**.

**Table S1: Classified taxa present in zooplankton groups.**

| **Zooplankton Groups** | | | | | |
|---|---|---|---|---|---|
| Cladocerans | Cyclopoids | Calanoids | Nauplia | Rotifers | Ciliates |
| *Bosmina sp.* | *Cyclops sp.* | *Eudiaptomus sp.* | Nauplius | *Asplanchna sp.* | *Paradileptus sp.* |
| *Daphnia sp.* | | | | *Kellicottia sp.* | *Askenasia sp.* |
| *Diaphanosoma sp.* | | | | *Synchaeta sp.* | *Coleps sp.* |
| *Daphnia cucullata* | | | | *Ascomorpha sp.* | *Didinium sp.* |
| *Chydorus sp.* | | | | *Keratella cochlearis* | *Strombidium sp.* |
| | | | | *Keratella quadrata* | Tintinnida |
| | | | | *Trichocerca sp.* | *Vorticella sp.* |
| | | | | *Conochilus sp.* | Ciliate unc1.1[1] |
| | | | | *Polyarthra sp.* | Ciliate uncl.2[1] |
| | | | | *Collotheca sp.* | Ciliate uncl.3[1] |
| | | | | Rotifer uncl.1[1] | Ciliate uncl.4[1] |
| | | | | Rotifer uncl.2[1] | Ciliate uncl.5[1] |
| | | | | Rotifer uncl.3[1] | Ciliate uncl.6[1] |

[1]*uncl. stands for unclassified*

### Table S2: Classified taxa present in phytoplankton groups.

| Phytoplankton Groups | | | | | |
|---|---|---|---|---|---|
| Chrysophytes | Cryptophytes | Chlorophytes | Diatoms | Cyanobacteria | Dinoflagellates |
| *Dinobryon sp.* | *Rhodomonas sp.* | *Closterium sp.* | *Asterionella sp.* | *Aphanizomenon sp.* | *Ceratium sp.* |
| *Uroglena sp.* | Cryptophyceae | *Coelastrum reticulatum* | *Fragilaria sp.* | *Limnoraphis sp.* | *Gymnodinium sp.* |
| *Synura sp.* | | *Cosmarium sp.* | *Aulacoseira sp.* | *Microcystis sp.* | *Peridinium sp.* |
| *Mallomonas akrokomos* | | *Gonium sp.* | *Synedra sp.* | *Coelosphaerium sp.* | Dinoflagellate uncl.1[1] |
| Dinobryon uncl.1[1] | | *Hormidium sp.* | *Synedra acus var. angustissima* | *Dolichospermum sp.* | |
| *Mallomonas sp.* | | *Tetraedron sp.* | Diatom unl.1[1] | *Snowella sp.* | |
| | | *Oocystaceae sp.* | Diatom uncl.2[1] | Chroococcales uncl.1[1] | |
| | | *Pandorina sp.* | Diatom uncl.3[1] | Chroococcales uncl.2[1] | |
| | | *Pediastrum sp.* | | Chroococcales uncl.3[1] | |
| | | *Phacotus sp.* | | Nostocales uncl.1[1] | |
| | | *Planktosphaeria sp.* | | | |
| | | *Scenedesmus sp.* | | | |
| | | *Staurastrum sp.* | | | |
| | | Chlorophyte uncl.1[1] | | | |
| | | Chlorophyte uncl.2[1] | | | |
| | | Chlorophyte uncl.3[1] | | | |
| | | Chlorophyte uncl.4[1] | | | |
| | | Chlorophyte uncl.5[1] | | | |
| | | Chlorophyte uncl.6[1] | | | |

[1]*uncl. stands for unclassified*

**Table S3: The performance of classifiers from the hierarchical image classification.**

| | Classifier | | |
|---|---|---|---|
| | Cyanobacteria | Phytoplankton | Zooplankton |
| **False Positives** | 0.002 | 0.214 | 0.038 |
| **Accuracy** | 0.986 | 0.907 | 0.946 |
| **F1 Score** | 0.882 | 0.863 | 0.934 |
| **AUC Macro** | 0.998 | 0.614 | 0.998 |
| **Precision** | 0.920 | 0.876 | 0.932 |
| **Recall** | 0.856 | 0.842 | 0.938 |

**Table S4: Performance of MDR S-maps from the best cross-validated parameters for each target cyanobacteria.**

| | RMVD S-maps Parameters | | | Cross-Validation[1] | | Fitting | | Persistence Model[2] | |
|---|---|---|---|---|---|---|---|---|---|
| | theta(θ)[3] | alpha(α)[4] | lambda(λ)[5] | RMSE | Rho | RMSE | Rho | RMSE | Rho |
| ***Microcystis group*** | 8 | 0.1 | 0.001 | 0.034 | 0.971 | 0.026 | 0.985 | 0.021 | 0.988 |
| ***Dolichospermum sp.*** | 8 | 0.1 | 0.001 | 0.030 | 0.952 | 0.023 | 0.975 | 0.016 | 0.985 |
| ***Chroococcus sp.*** | 8 | 0.1 | 0.001 | 0.038 | 0.962 | 0.029 | 0.981 | 0.023 | 0.986 |
| ***Coelosphaerium sp.*** | 8 | 0.1 | 0.001 | 0.036 | 0.965 | 0.028 | 0.982 | 0.022 | 0.986 |

[1]*Sequential leave-future-out cross-validation*

[2]*Persistence model at a 1 day prediction horizon*

[3]*S-maps localization parameter determining the degree of nonlinearity (28)*

[4]*Elastic net mixing parameter between Lasso and Ridge pendalty (37)*

[5]*Elastic net complexity parameter (37)*

**Table S5: The performance of Random Forest models for weekly predictions of focal nutrients.**

| | Cross-Validation | | | | Testing | |
|---|---|---|---|---|---|---|
| | Mean RMSE | SD RMSE | Mean $R^2$ | SD $R^2$ | RMSE | $R^2$ |
| **Ammonium (μg/L)** | 37.974 | 13.531 | 0.678 | 0.248 | 43.536 | 0.551 |
| **Nitrate (mg/L)** | 0.211 | 0.047 | 0.670 | 0.143 | 0.166 | 0.745 |
| **Phosphate (μg/L)** | 4.801 | 1.616 | 0.854 | 0.120 | 6.328 | 0.600 |

**Table S6: Blooming point change across gradients of water temperature of phosphorus levels for target biotic factors and cyanobacterial taxon.** These changes are calculated by varying biotic factors from their minimum to their maximum observed values, and taking the resulting variation in temperature and $PO_4$ levels.

| | | Blooming Point Change[2] | | | |
|---|---|---|---|---|---|
| **Biotic Factors** | **Abiotic condition** | **Microcystis** | **Dolichospermum** | **Chroococcus** | **Coelosphaerium** |
| Cladocerans | Temperature | - 10.123 | - 0.913 | + 5.061 | - 2.323 |
| Calanoids | Temperature | - 4.149 | + 8.38 | + 1.328 | + 4.895 |
| Rotifers | Temperature | na[1] | + 13.442 | - 3.07 | - 0.996 |
| Diatoms | Temperature | na[1] | + 1.742 | + 1.577 | + 1.659 |
| Chrysophyta | Temperature | - 8.131 | - 8.629 | - 3.07 | + 0.996 |
| Cyanobacteria | Temperature | na[1] | + 10.455 | - 0.747 | - 4.481 |
| Cladocerans | $PO_4$ | - 14.523 | - 8.684 | + 6.139 | - 4.791 |
| Calanoids | $PO_4$ | - 18.416 | + 18.565 | + 2.695 | + 9.133 |
| Rotifers | $PO_4$ | + 7.336 | + 20.661 | - 1.647 | - 0.299 |
| Diatoms | $PO_4$ | - 18.266 | + 3.444 | + 1.048 | + 3.144 |
| Chrysophyta | $PO_4$ | - 27.548 | - 29.495 | - 14.074 | + 1.198 |
| Cyanobacteria | $PO_4$ | - 16.02 | + 9.882 | + 2.246 | - 7.486 |

*[1]Cases when cyanobacterial ngGR>0 or <0*

*[2]Bloomings point is the intercept of the zero net growth isocline (nGR=0)*

*Changes in blooming point are in °C for temperature and µg/L for $PO_4$*

*+/- indicate increase/decrease in blooming requirements*
